# gr Predictor: a Deep-Learning Model for Predicting the Hydration Structures around Proteins

**DOI:** 10.1101/2022.04.18.488616

**Authors:** Kosuke Kawama, Yusaku Fukushima, Mitsunori Ikeguchi, Masateru Ohta, Takashi Yoshidome

**Author notes:** **Corresponding Author Takashi Yoshidome** - Department of Applied Physics, Graduate School of Engineering, Tohoku University, Sendai 980-8579, Japan;.

## Abstract

Among the factors affecting biological processes such as protein folding and ligand binding, hydration, which is represented by a three-dimensional water-site-distribution-function around the protein, is crucial. The typical methods for computing the distribution functions, including molecular dynamics simulations and the three-dimensional reference interaction site model (3D-RISM) theory, require a long computation time from hours to tens of hours. Here, we propose a deep-learning model rapidly estimating the distribution functions around proteins obtained by the 3D-RISM theory from the protein 3D structure. The distribution functions predicted using our deep-learning model are in good agreement with those obtained by the 3D-RISM theory. Particularly, the coefficient of determination between the distribution function obtained by the deep-learning model and that obtained using the 3D-RISM theory is approximately 0.98. Furthermore, using a graphics processing unit (GPU), the calculation by the deep learning model is completed in less than one minute, more than 2 orders of magnitude faster than the calculation time of 3D-RISM theory. Therefore, our deep learning model provides a practical and efficient way to calculate the three-dimensional water-site-distribution-functions. The program called “gr Predictor” is available under the GNU General Public License from https://github.com/YoshidomeGroup-Hydration/gr-predictor.

**Table of Contents graphic:** 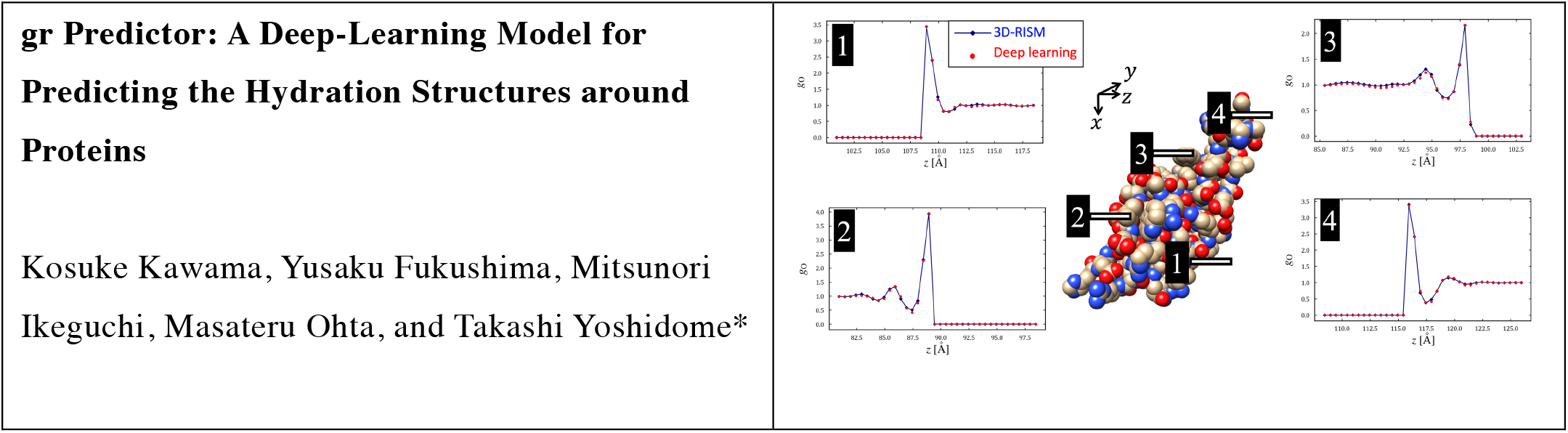

## 1. INTRODUCTION

Protein hydration is one of the factors governing the biophysical processes involving the protein, including folding and ligand binding^1,2^. Particularly, hydration strongly affects the stability of native structure and denaturation of proteins, whereas the ligand-protein complex is often stabilized by water-mediated interactions between the ligand and protein. Thus, elucidating the protein hydration properties is crucial to understand the biophysical processes and perform structure-based drug design in consideration of water molecules.

Hydration of protein is characterized by the three-dimensional water-site-distribution-functions around the protein. The distribution functions can be obtained using molecular dynamics (MD) simulations and the three-dimensional reference-interaction site model (3D-RISM) theory^3^. MD simulations exactly compute the three-dimensional water site distribution functions, whereas the 3D-RISM theory, which is a statistical mechanical theory of solvation, approximately computes the distribution functions with the force fields employed by the MD simulations. The usefulness of the distribution functions has been demonstrated. As an example, the performance of the deep-learning (DL) model for the pose prediction improved upon the incorporation of the three-dimensional water-site-distribution-functions, obtained with MD simulations, inside the ligand-bind pocket^4^. The distribution function obtained using the 3D-RISM theory has also been widely discussed. The position of the crystal waters inside the cavity of hen egg-white lysozyme is difficult to compute via MD simulation, because the movement of a water molecule from outside the protein towards inside is hardly attained in a reasonable simulation time; conversely, the three-dimensional water-site-distribution-functions obtained with the 3D-RISM theory successfully reproduced the crystal-water positions^5^. Furthermore, the partial molar volume (PMV) that can be computed with the Kirkwood-Buff solution theory combined with the three-dimensional water-site-distribution-functions^6^ exhibited a perfect agreement with the experimental data of several proteins using the distribution functions obtained with the 3D-RISM theory^7,8^.

The advantages of the 3D-RISM theory over the MD simulations include the shorter computation time for obtaining the three-dimensional water-site-distribution-functions. Exploiting the power of a supercomputer and the advantage of the shorter computation time of the 3D-RISM theory, we have statistically analyzed the hydration states of 3,706 static crystallographic structures of a protein^9^. However, the investigation of amino acid mutations, ligand binding processes, and protein conformational changes require more than hundred thousand of protein structures. To this aim, the 3D-RISM theory is inadequate, because the calculation of the hydration state of a protein requires a few hours with a single central processing unit (CPU). Thus, a new method drastically reducing the computation time of the 3D-RISM theory should be developed.

In the present paper, we propose a DL model for predicting the three-dimensional water-site-distribution-functions around the proteins obtained with the 3D-RISM theory. The data used for training the DL model, network architecture, prediction accuracy, and computation time of the DL model are described. Finally, a comparison of our DL model with other methodologies for obtaining the hydration states around proteins is discussed. Because our DL model accurately reproduced the distribution functions obtained with the 3D-RISM theory with a computation time of less than one minute using a single graphics processing unit (GPU), our DL model enables us to investigate amino acid mutations, ligand binding processes, and protein conformational changes.

## 2. MATERIALS AND METHODS

### Proteins used for the computation

Twenty-seven proteins were selected from the proteins used in our previous study^9^ considering 3,706 proteins taken from the protein-ligand complexes deposited in the PDBbind refined set (v. 2017)^11-16^. The selection for this study was performed as follows. From the initial 3,706 proteins, only the proteins without ions were considered, to reduce the number of atom types in the deep-learning model. This led to the selection of 2,718 proteins summarized in “Data2718-SI-Forsubmit.xlsx”. Afterwards, the twenty-seven proteins shown in Table 1 were randomly selected. Among the twenty-seven proteins, twenty-two proteins were used for training, and the remaining five proteins were used for the test. The following preprocesses were conducted to the 3,706 proteins in the previous study^9^: the ligand and the crystal waters were removed, and the chain closest to the ligand was employed for the protein structure with multiple chains.

**Table 1.** Proteins used for developing and evaluating the deep-learning model. “Train: 10” and “Train: 12” are equal to “Train: Ten proteins” and “Train: Twelve proteins”, respectively.

| PDB ID | Structure Title | High Reso. Limit (Å) | Dataset |
| --- | --- | --- | --- |
| 1A30 | HIV-1 protease complexed with a tripeptide inhibitor | 2.00 | Train: Ten proteins |
| 1FCH | PTS1 complexed to the TPR region of human PEX5 | 2.20 | Train: 10 |
| 1PZ5 | Antibody in complex with octapeptide | 1.80 | Train: 10 |
| 2CE9 | A peptide bound to the Groucho-TLE WD40 domain. | 2.12 | Train: 10 |
| 2HKF | The complex Fab M75- Peptide | 2.01 | Train: 10 |
| 2PV1 | SurA complexed with peptide WEYIPNV | 1.30 | Train: 10 |
| 2QBW | PDZ-Fibronectin fusion protein | 1.80 | Train: 10 |
| 3BZF | Major histocompatibility in complex with HLA-E | 2.50 | Train: 10 |
| 3DRF | OppA complexed with an endogenous peptide | 1.30 | Train: 10 |
| 3DRI | OppA co-crystallized with an octamer peptide | 1.80 | Train: 10 |
| 3ERY | H-2 class I histocompatibility antigen in complex with peptide | 1.95 | Train: Twelve proteins |
| 3G19 | ClpS protease adaptor protein in complex with peptide | 1.85 | Train: 12 |
| 3IFL | Amyloid beta peptide:antibody complex | 1.50 | Train: 12 |
| 3P9M | H2-Kb in complex with epitope OVA-G4 | 2.00 | Train: 12 |
| 3T6B | human DPPIII in complex with Tynorphin | 2.40 | Train: 12 |
| 3TCG | E. coli OppA complexed with the tripeptide KGE | 2.00 | Train: 12 |
| 3UPV | pHsp70-complex of yeast Sti1 | 1.60 | Train: 12 |
| 4EZR | E.coli DnaK in complex with drosocin | 1.90 | Train: 12 |
| 4EZZ | E.coli DnaK in complex with peptide ELPLVKI | 2.05 | Train: 12 |
| 4YNL | HetR in complex with the hexapeptide<br>ERGSGR | 2.10 | Train: 12 |
| 5E6O | C. elegans LGG-2 bound to an AIM/LIR<br>motif | 1.80 | Train: 12 |
| 5LSO | SPF45 UHM domain with cyclic peptide<br>inhibitor | 2.22 | Train: 12 |
| 2HA2 | Acetylcholinesterase complexed with<br>succinylcholine | 2.05 | Test |
| 2O4L | HIV-1 Protease in Complex with Tipranavir | 1.33 | Test |
| 3JVR | Chk1 complexed with allosteric inhibitor | 1.76 | Test |
| 3O5N | Shank PDZ domain complexed with small<br>molecule | 1.83 | Test |
| 4KAO | Focal adhesion kinase in complex with<br>inhibitor | 2.39 | Test |

The similarities of the sequences between the twenty-seven proteins are discussed in text S1 (Supporting Information). None of the proteins used in the test had 90% sequence similarity to the twenty-two proteins used in the training. The effects of the random selection of the twenty-seven proteins are discussed in the subsection “Discussion of the selection of twenty-seven proteins”.

### 3D-RISM theory

In our previous study^9^, the 3D-RISM theory was applied to obtain the distribution function at the position ***r*** for the water site *α* = H (hydrogen) or O (oxygen), denoted by *g*_*α*_(***r***) hereinafter. In the present study, *g*_*α*_(***r***) was used as a target variable for the construction of the DL model. In the following, the force fields and parameters employed in the previous study^9^ are described. The distribution functions were obtained using the Amber ff99SB force-fields^17^ for the proteins, whereas the coincident SPC/E model^18^ was employed for the water molecule. The values of the dielectric constant, bulk density, and temperature were 78.497, 0.03332 Å^-3^, and 310 K, respectively. In the computation using the 3D-RISM theory, a water box surrounding the protein was prepared so that the minimum distance between the protein and the edge of the box was 14 Å. The linear grid spacing of 0.5 Å was set for the *x, y*, and *z* coordinates.

### Input and output formats for the deep-leaning model

To input a protein structure into our DL model, the protein structure was converted into the voxel format schematically illustrated in Fig. 1. First, the protein was decomposed into five atom types composed of carbon, nitrogen, oxygen, sulfur, and hydrogen. Afterwards, the box surrounding the protein with the same size as that of the water box used for the 3D-RISM theory was prepared. The grid size of the voxel and the position of the protein were also the same as those used for the computation using the 3D-RISM theory. Then, the contribution of the *i*-th atom of atom type *j* to *k*-th voxel, *n*(*k, i, j*), was computed in accordance with Eq. (1)^19^:

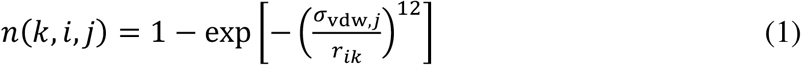

where *σ*_vdw,*j*_ is the van-der-Waals-radius of the atom type *j*, and *r*_*ik*,_ is the distance between the *i*-th atom and the position of the *k*-th voxel. The value of *σ*_vdw,*j*_ for each atom type is summarized in Table S1 in the Supporting Information. Finally, the contribution of the atom type *j* to *k*-th voxel, *n*(*k, i, j*), *N*(*k, j*), was computed according to Eq. (2):

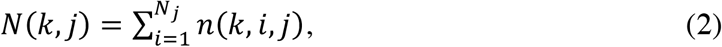

where *N*_*j*_ is the number of atoms for the atom type *j* in the protein. Through this procedure, the protein structure was decomposed into five boxes according to the atom type, with each box composed of the voxels with the value of *N*(*k, j*).

**Fig. 1.**
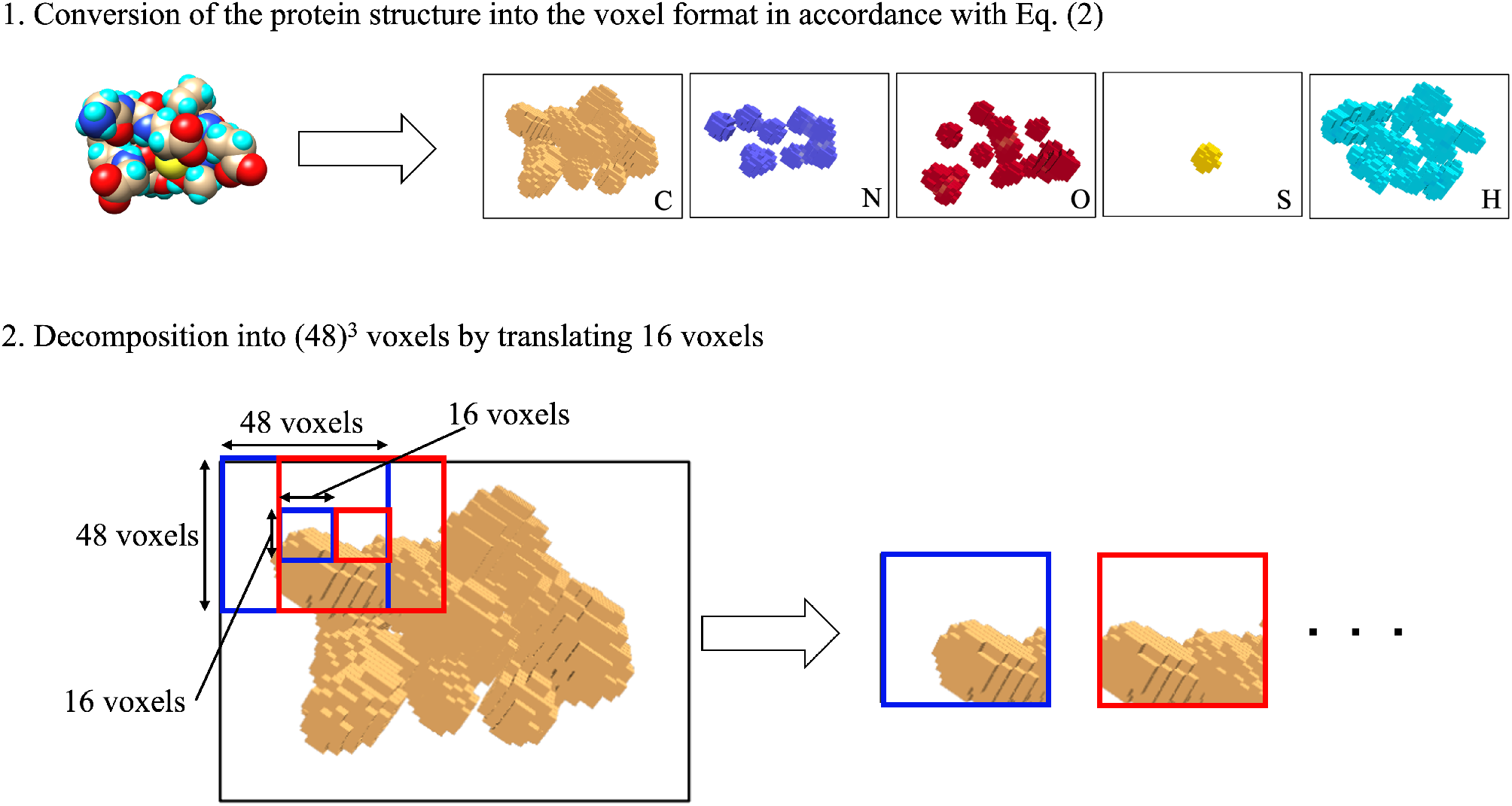
Schematic of the conversion of a protein structure into the voxel format.

By decomposing each box into small boxes of 48^3^ voxels (Fig. 1), we made our DL model applicable to proteins with arbitrary sizes. Hereinafter, the box is referred to as the “partial protein box” and the set of the boxes of five atom types at the same position in the protein is referred to as the “set of partial protein box”. To exclude the effect of the boundary of the partial protein box on the training and test by conducting them with the central 16^3^ voxels, the decomposition was performed by translating the partial protein box by 16 voxels (Fig. 1).

The output format of our DL model was the same as that of the water box of *g*_*α*_(***r***) (*α* = H or O) obtained using the 3D-RISM theory. For the training of our DL model, the water box was also decomposed into the boxes of 48^3^ voxels as previously described. The resulting box is referred to as the “partial water box”. With a set of partial protein box as input, our DL model outputs the corresponding partial water box. By summing the central 16^3^ voxels of each partial water box, *g*_H_(***r***) or *g*_O_(***r***) were obtained.

### Deep-learning model

For the network architecture for our DL model that predicts the protein hydration structure, we employed the U-net^20^. As schematically shown in Fig. 2, the U-net is an encoder-decoder type architecture^21^. The deep-learning model was constructed for predicting a distribution function *g*_*α*_(***r***) (*α* = H or O).

**Fig. 2.**
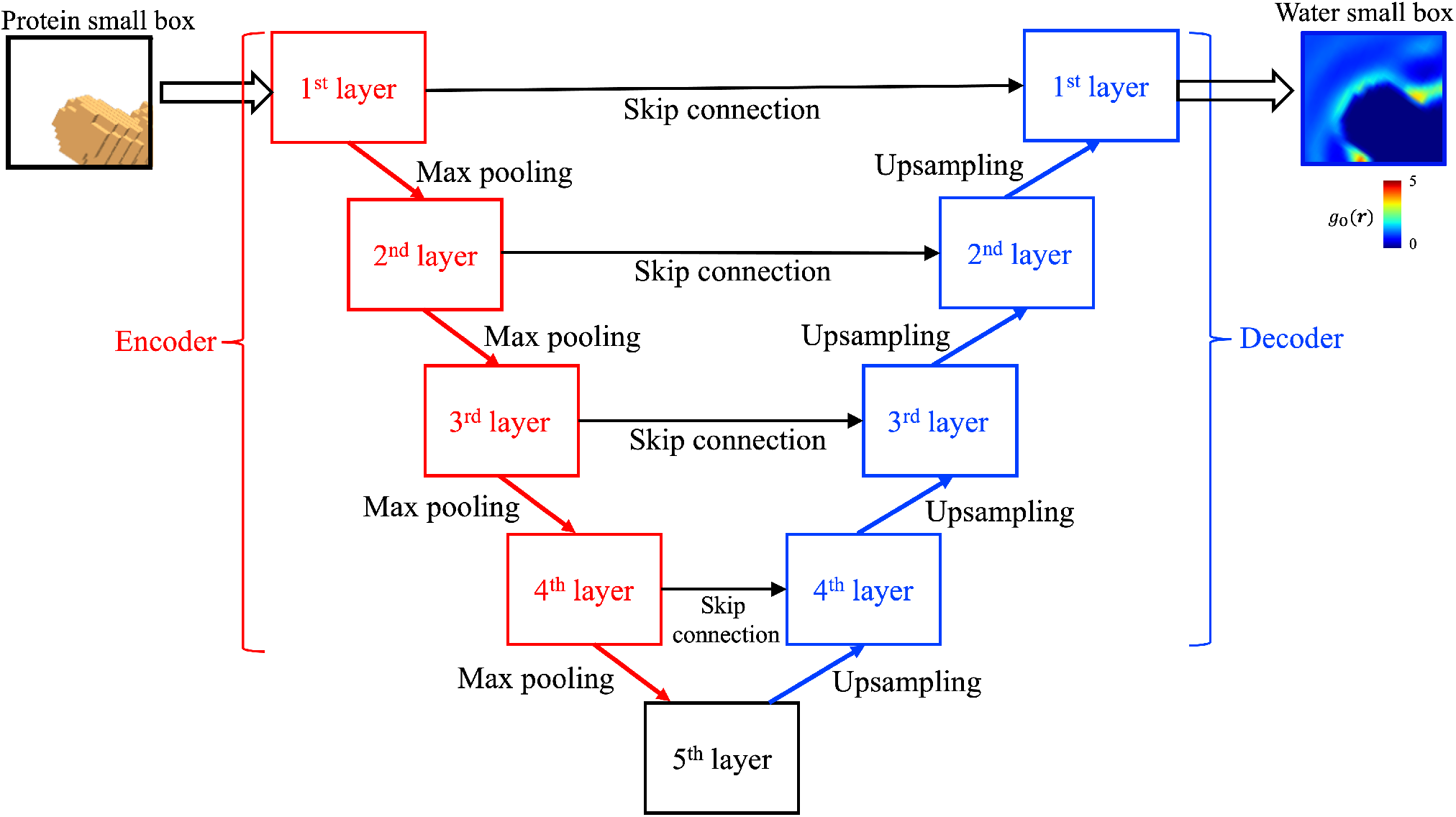
Schematic of the U-net architecture.

The architecture we employed was essentially the same as that used in the original U-net model used for biomedical image segmentation^10^. As shown in Fig. 2, both encoder and decoder consisted of four layers, referred to as the “encoder-decoder layers”, and a 5^th^ layer was prepared between the 4^th^ layers of encoder and decoder. Each encoder-decoder layer consisted of two convolutional layers, each followed by the activation using the ReLU function. A 2×2×2 max-pooling layer was added after the second convolutional layer of the encoder-decoder layer in the encoder. The max-pooling layer was replaced by an upsampling layer^20^ in the encoder-decoder layer in the decoders. The number of filters in the first convolutional layers in an encoder-decoder layer was doubled from the previous layer in the encoder, and a half from the previous layer in the decoder, respectively. Skip connection was added according to the original U-net architecture.

Our DL model differed from the original U-net model in four aspects. First, while the original U-net model was implemented for two-dimensional images, our model was implemented for three-dimensional data of a partial protein box and a partial water box. Furthermore, in our model the zero-padding was added in the convolutional layers. Moreover, when training our DL model, the dropout layer was also added after the ReLU activation in the first convolutional layer, to reduce the overfitting. Finally, the convolution followed by the up-sampling in the original U-net architecture was removed in our architecture. The effect of convolution in the upsampling layer was small (text S2 in the Supporting Information).

As shown in Table 2, our DL model had four hyperparameters, collectively referred to as “hyperparameter set”, one of which was the filter size (i.e., number of voxels in the filters) in the convolutional layers. Three sizes were prepared, namely 3^3^, 4^3^, and 5^3^. The number of filters (*N*_Firstfilter_) for an atom type at the first encoder-decoder layer in the encoder was another hyperparameter. Because the number of atom types was five, the total number of filters was 5·*N*_Firstfilter_. The third hyperparameter was the number of voxels whose value was set at zero in the dropout layer, *N*_D_. The ratio of *N*_D_ and the total number of voxels in the dropout layer, referred to as “dropout ratio”, was set to 0.3 or 0.5. Finally, the dropout was applied to the 5^th^ layer, 4^th^–5^th^ layers, 3^rd^–5^th^ layers, 2^nd^–5^th^ layers, or all layers for each of (i) only the encoder, (ii) only the decoder, or (iii) both the encoder and the decoder. The case in which no dropout was applied to both the encoder and the decoder was also considered.

**Table 2.** Hyperparameters and optimization ranges of our deep-learning model.

| Hyperparameter | Range of the parameters for optimization |
| --- | --- |
| A. The size of filter for convolution | $[3^3]$, $[4^3]$, or $[5^3]$ |
| B. The number of filters at the first layer in the encoder | 16 or 32 |
| C. The dropout ratio | 0.3 or 0.5 |
| D. The layers to which the dropout is applied | ✓ The dropout was applied to (i) only the encoder, (ii) only the decoder, or (iii) both the encoder and the decoder.<br>✓ For each of (i), (ii), and (iii), the dropout was applied to 5<sup>th</sup> layer, 4<sup>th</sup>-5<sup>th</sup> layers, 3<sup>rd</sup>-5<sup>th</sup> layers, 2<sup>nd</sup>-5<sup>th</sup> layers, or all layers.<br>✓ The case in which no dropout was applied to both the encoder and the decoder was also considered. |

Our DL model was implemented using the TensorFlow library (2.1.0) for the models predicting *g*_*α*_(***r***) (*α* = H or O). The training was performed using Adam optimizer with default parameters.

### Hyperparameter optimization

The hyperparameters were optimized through the two-fold cross validation. The twenty-two proteins used for the training were split into two sets to homogeneously adjust the total number of partial protein boxes of two sets: ten proteins and the remaining twelve proteins (Table 1). The number of partial protein boxes (*N*_TData_) was 6,858 and 7,101 from the ten and twelve proteins, respectively. The number of partial water boxes was also *N*_TData_. In the cross validation, when the ten proteins were used for the training of our DL model, the remaining twelve proteins were used for the validation of the model and vice versa. Hereinafter, the data for the training and validation are denoted by “training data” and “validation data”, respectively. Each data is composed of the partial protein boxes and the corresponding partial water boxes.

For a hyperparameter set of our DL model, the number of epochs was set to 200 and the mean-square error in Eq. (3) was employed for the loss function, *E*:

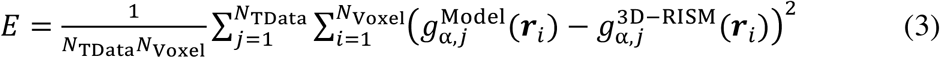

where *N*_Voxel_ is the number of voxels in the box, equal to 16^3^; 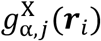 (X=“Model” or “3D-RISM”, and α=“O” or “H”) is the *g*_*α*_(***r***) value at the voxel position of ***r***_*i*_ for the *j*-th box; the superscripts “Model” and “3D-RISM” indicate the *g*_*α*_(***r***) obtained from our DL model and the 3D-RISM theory, respectively. Hereinafter, the *E* value at *i* epoch is denoted by *E*_Train_(*i*).

For each epoch (*i*) in the training, we also computed the loss function of Eq. (3) using the validation data (*E*_Validation_(*i*)) to check whether the overfitting did not occur during the training. In the computation, all dropout layers used in the training were not used. The possible overfitting was checked by the comparability of the *E*_Validation_(200) and *E*_Training_(200) values.

Each of the four hyperparameters was set to a value within the range shown in Table 3, leading to 162 hyperparameter sets. For each hyperparameter set, the following computations were performed: (1) Training was performed with the ten proteins and the corresponding *g*_H_(***r***) or *g*_O_(***r***) as the training data; (2) the *E*_Validation_(200) value was saved; (3) the procedures (1) and (2) were repeated with the twelve proteins for the training, and (4) the average of the two *E*_Validation_(200) values, namely *Ē*_Validation_(200), was computed. After the computations for all the 162 hyperparameter sets, the hyperparameter set with the smallest *Ē*_Validation_(200) value, denoted by “optimized hyperparameter set”, was selected.

**Table 3.** The optimized hyperparameter and statics of our deep-learning model. The *E*_Validation_(200) values obtained using the ten and twelve proteins are denoted as 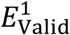 and 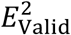, respectively.

|  |  |  |
| --- | --- | --- |
| Hyperparameter | A. The size of filter for convolution | $[3^3]$ |
|  | B. The number of filters at the first layer in the encoder | 32 |
|  | C. The dropout ratio | 0.3 |
|  | D. The layers to which the dropout is applied | ✓ both encoder and decoder<br>✓ 2<sup>nd</sup>-5<sup>th</sup> layers |
| Statistics | $E_{\text{Valid}}^1$ | 0.0045 |
| | $E_{\text{Valid}}^2$ | 0.0039 |
| | Average of $E_{\text{Valid}}^1$ and $E_{\text{Valid}}^2$ | 0.0042 |

### Tests

After the training using the optimized hyperparameter set and the twenty-two proteins, *g*_*α*_(***r***) (α=“O” or “H”) for the five proteins described in Table 1 was computed as a test. The training procedure was the same as that described in the previous subsection. In the test, each protein was first converted into the voxel format described in “Input and output formats for the DL model” subsection. Afterwards, using a partial protein box, *g*_*α*_(***r***) of the corresponding partial water box was predicted using our DL model. Finally, *g*_*α*_(***r***) of the whole protein was obtained by summing the central 16^3^ voxels of the predicted water boxes.

To quantitatively compare the peak positions of *g*_O_(***r***) of our DL model with those of the 3D-RISM theory, the following analysis was conducted. First, water oxygen atoms were placed using the program Placevent^22^, in which water oxygen atoms were placed using the *g*_O_(***r***) values. In the program, the placement of water oxygen atoms was performed in three steps: (i) A water oxygen atom was placed at the position of the voxel with the largest *g*_O_(***r***) value (denoted by ***r***_Max_); (ii) The region ***δ*** satisfying Eq. (4) was identified:

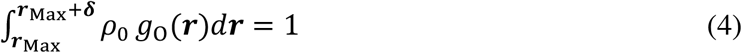

(iii) The *g*_O_(***r***) values at the voxels within the region ***δ*** were set at zero (the obtained *g*_O_(***r***) is referred to as “new *g*_O_(***r***)”); (iv) Steps (i), (ii), and (iii) were repeated using the new *g*_O_(***r***) until *g*_O_(***r***_Max_) < 1.5, which is the default value in the program Placevent. The placement of water oxygen atoms was performed using the *g*_O_(***r***) values obtained with the 3D-RISM theory and those obtained with our DL model. The positions of the *i*-th water oxygen atom obtained using the 3D-RISM theory and that using our DL model are denoted by 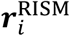 and 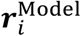, respectively. The number of placed water oxygen atoms is denoted by *N*_RISM_ and *N*_Model_, and typically *N*_RISM_ ≠ *N*_Model_ because *g*_O_(***r***) obtained using our DL model was slightly different from that obtained using the 3D-RISM theory.

Afterwards, for each water oxygen atom placed using *g*_O_(***r***) obtained with the 3D-RISM theory, the distance *D*_*i*_ defined in Eq. (5) was computed:

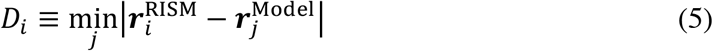

The average of *D*_*i*_ among the water oxygen atoms placed using the 3D-RISM theory and its standard deviation were obtained to analyze the results and histogram of *D*_*i*_.

To investigate the prediction performance at the ligand-binding pocket from the viewpoint of *D*_*i*_, the following analysis was further performed. The water oxygen atoms at the ligand-binding pocket were defined by the placed water oxygen atoms within 5 Å from the heavy atoms of the ligand. For each water oxygen atom placed using the *g*_O_(***r***) of 3D-RISM theory, the *D*_*i*_ value was computed.

Finally, the prediction performance from the viewpoint of the positions of crystal waters was studied via the analysis of the crystal waters within 5 Å from the heavy atoms in the protein. For each crystal water, *D*_*i*_ was computed.

## 3. RESULTS AND DISCUSSION

In this section, the results obtained for *g*_O_(***r***) are presented, whereas those *g*_H_(***r***), which were analogous to those for *g*_O_(***r***), are discussed in the subsection “Prediction of the distribution function of water hydrogen site”.

### Cross validations

To determine the optimized hyperparameter set, a two-fold cross validation was conducted as described in “Optimization of the hyperparameters”. The number of partial protein boxes was 6,858 and 7,101 from the ten and twelve protein sets, respectively. The error values at 200 epoch defined by Eq. (3) were computed using the ten and twelve proteins, and their *E*_Validation_(200) and *Ē*_Validation_(200) values are summarized in Table S1.

The optimized hyperparameter with the smallest *Ē*_Validation_(200) value was the hyperparameter set number 44 (Table S2). The hyperparameter values and their statistics are summarized in Table 3. The *Ē*_Validation_(200) value, namely the difference between the *g*_O_(***r***) values of 3D-RISM and of our DL model, was sufficiently small (0.0042). The corresponding average deviation of *g*_O_(***r***) obtained by our DL model from *g*_O_(***r***) obtained by the 3D-RISM theory was 0.06. As shown in Fig. S1, *Ē*_Validation_(200) was analogous to *Ē*_Train_ (200), indicating the absence of overfitting.

The results reported in the next paragraphs were performed using the DL model trained with the optimized hyperparameter set. The results for the other hyperparameters are discussed in text S3 (Supporting Information).

### Prediction tests

The correlation between the *g*_O_(***r***) values predicted by our DL model and those calculated by the 3D-RISM theory, coefficient of determination R^2^ score values, and root mean square error (RMSE) of five proteins for the test are shown in Fig. 3. For the five proteins tested, the R^2^ values were high and most of the points resided close to the line representing *y*=*x*. Moreover, the RMSE values indicated the accuracy of the *g*_O_(***r***) prediction of our DL model. The encouraging result on the accuracy was accompanied by a drastic decrease of computation time of two orders of magnitude: the computation was completed within a minute with our DL model and a single GPU.

**Fig. 3.**
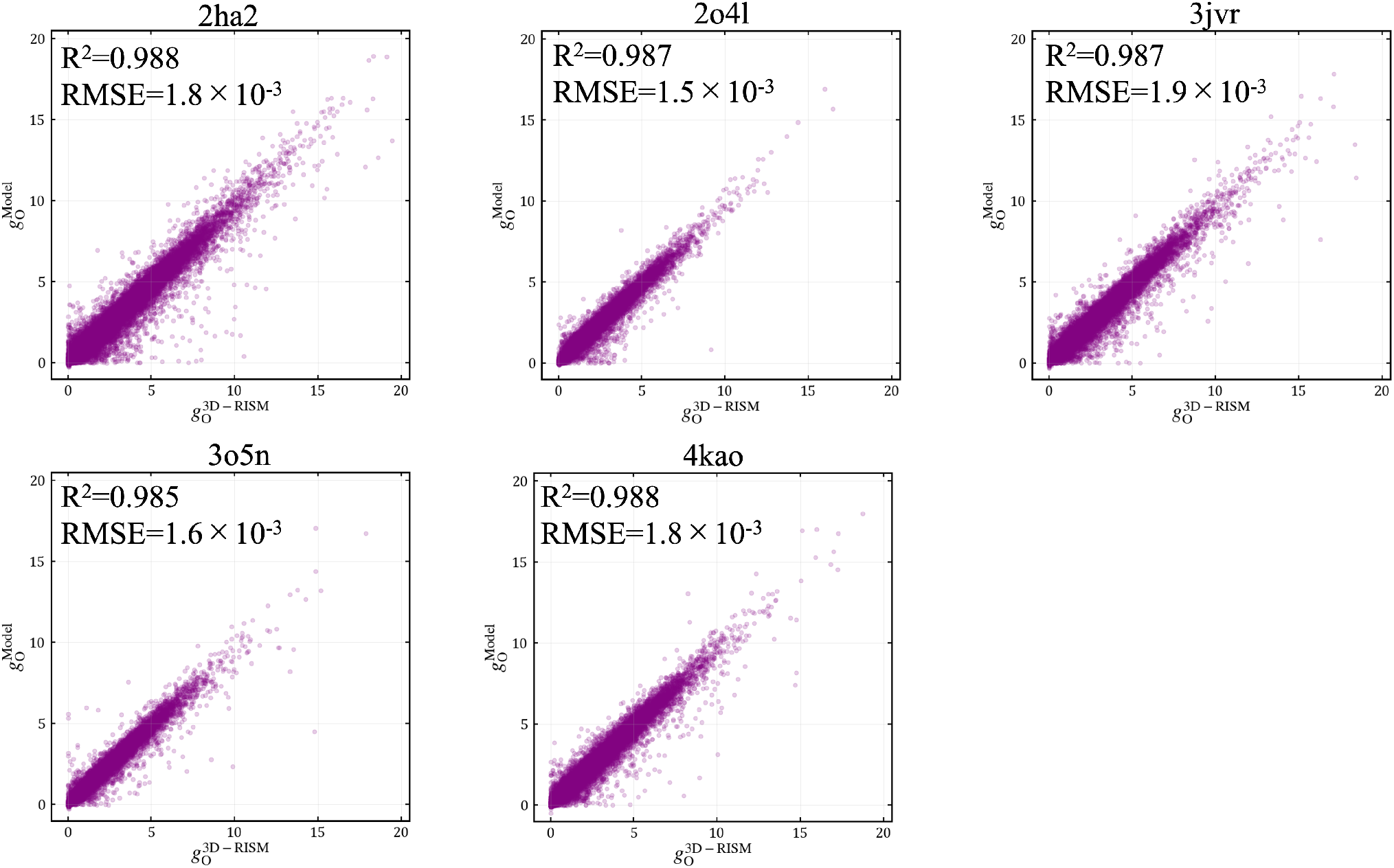
Correlation between the *g*_O_(***r***) values predicted by our deep-learning model and those calculated by the 3D-RISM theory. The coefficient of determination R^2^-score values and root mean square error (RMSE) are indicated.

Furthermore, the comparison was performed with the voxels in the ligand-binding pocket defined by those within 5 Å from the heavy atoms in the ligand (Fig. 4). The prediction performance of our DL model was high also in the ligand-binding pocket: most of the points resided close to the line *y*=*x*, with high R^2^ values. Therefore, our DL model can successfully predict the hydration structure in the ligand-binding pocket. High prediction-accuracy of *g*_O_(***r***) in the ligand-binding site is important for the structure-based design of new molecules using the information of the hydration in the binding site.

**Fig. 4.**
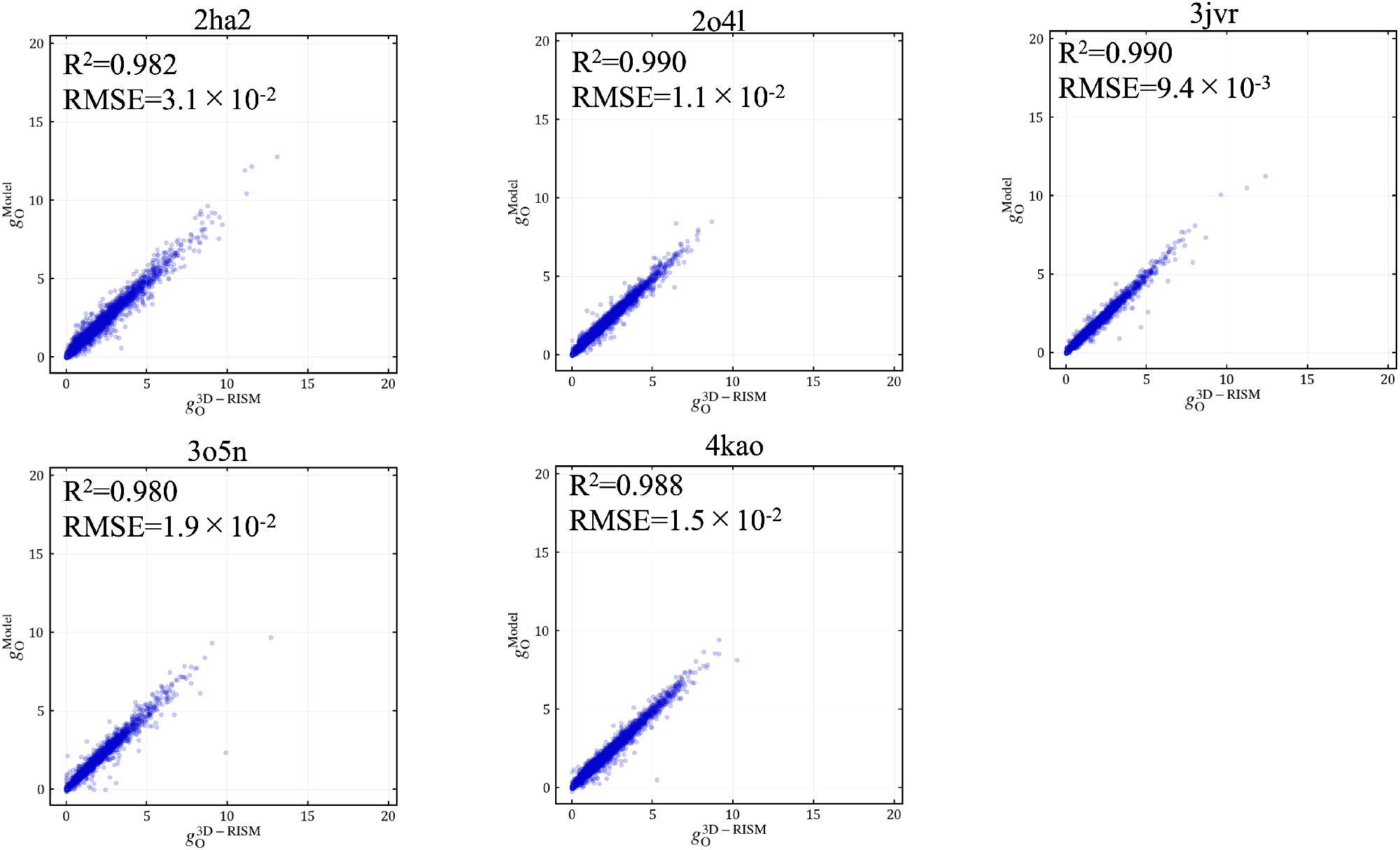
Results at the ligand-binding site for the five test proteins. Correlation between the *g*_O_(***r***) values predicted by our deep-learning model and those calculated by 3D-RISM theory. The coefficient of determination R^2^-score values and root mean square error (RMSE) are indicated.

For all the proteins, the RMSE value of the ligand-binding site was larger than that of all points, reasonably due to the value of *g*_O_(***r***) at the bulk region. It was found from slice 5 of Fig. 5 that the prediction performance at the bulk region is good: *g*_O_(***r***)at the bulk region was one for both of the 3D-RISM theory and our DL model. This result explains the RMSE value of all points, including most of the bulk points, smaller than that of the ligand-binding site.

**Fig. 5.**
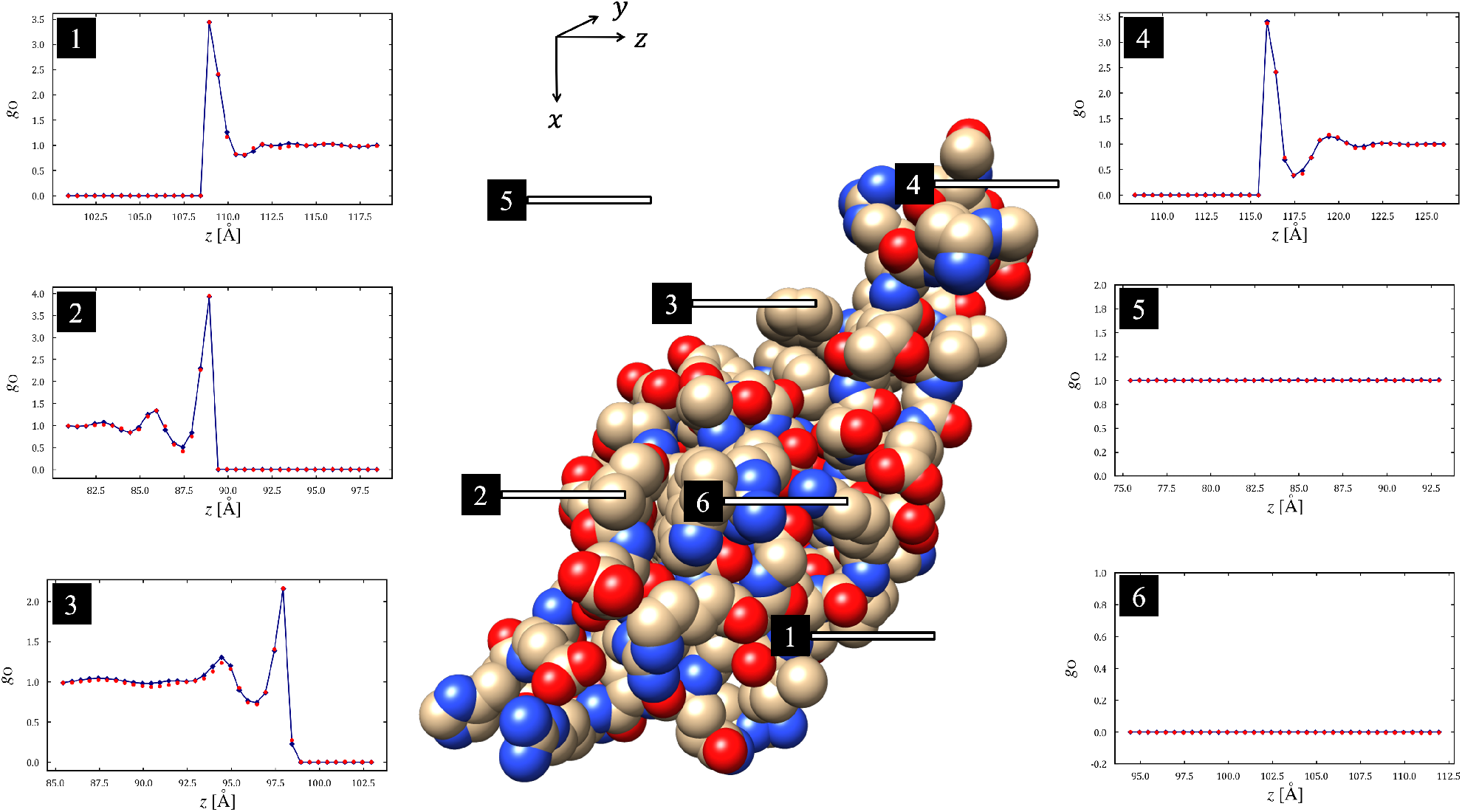
Results of *g*_O_(***r***) for shank3 PDZ domain (PDB code: 3o5n). The *g*_O_(***r***) values at the six-line regions illustrated in the protein are shown. The blue lines and blue points represent the *g*_O_(***r***) values obtained using the 3D-RISM theory, whereas the red points represent the *g*_O_(***r***) values obtained with our deep-learning model.

Finally, the results for shank3 PDZ domain (Protein Data Bank (PDB) code: 3o5n) are shown in Fig. 5 to discuss how our DL model reproduced the *g*_O_(***r***) values in detail. The agreement between the *g*_O_(***r***) values of the 3D-RISM theory and those of our DL model was good, as both the peak heights and the peak positions were well predicted by our DL model. Additionally, our DL model reproduced the *g*_O_(***r***) values inside the protein (slice 6 in Fig. 5) and those at a bulk region (slice 5 in Fig. 5). Moreover, a high R^2^-score value (0.985) indicated the good correlation between *g*_O_(***r***) values of our DL model and those of 3D-RISM theory (Fig. 3). However, for few points, the *g*_O_(***r***) values of our DL model deviated from those of the 3D-RISM theory. Particularly, the points with large deviation corresponded to the areas in the cavities with a size comparable to that of the water molecule (Fig. S2). The current training data did not contain sufficient data for such cavities. Adding such data would therefore improve the performance of our DL model.

### Placement of water oxygen atoms

To discuss how our DL model successfully predicted the peak positions of *g*_O_(***r***), water oxygen atoms were placed at the *g*_O_(***r***) peaks using the program Placevent. For the placement of water oxygen atoms, the values of *g*_O_(***r***) obtained using the 3D-RISM theory and our DL model were used. The histograms of *D*_*i*_ for the five proteins are shown in Fig. 6, whereas the average of *D*_*i*_, *N*_RISM_, and *N*_Model_ for each protein are summarized in Table 4. The histograms related to the water molecule placed at the point *g*_O_(***r***)>1.5 (probability 1.5 times higher than that of a bulk water) in Fig. 6 (a), (d), (g), (j), and (m) indicate that approximately 60% of the water oxygen atoms of our DL model was placed within 0.5 Å from the water oxygen atoms of the 3D-RISM theory. The average of *D*_*i*_ was 0.6–0.7 Å for all five proteins (Table 4 and Fig. 7). The calculated value was 1/4–1/5 of the Lennard-Jones sigma value, associated to the radius of the atom, of the water oxygen atom for the coincident SPC/E model (3.17 Å). Therefore, the *g*_O_(***r***) peak positions obtained using our DL model were close to those obtained using the 3D-RISM theory. Essentially the same results were obtained for the *g*_O_(***r***) values at the ligand-binding pocket (Fig. 6(b), (e), (h), (k), (n), and Table 5).

**Fig. 6.**
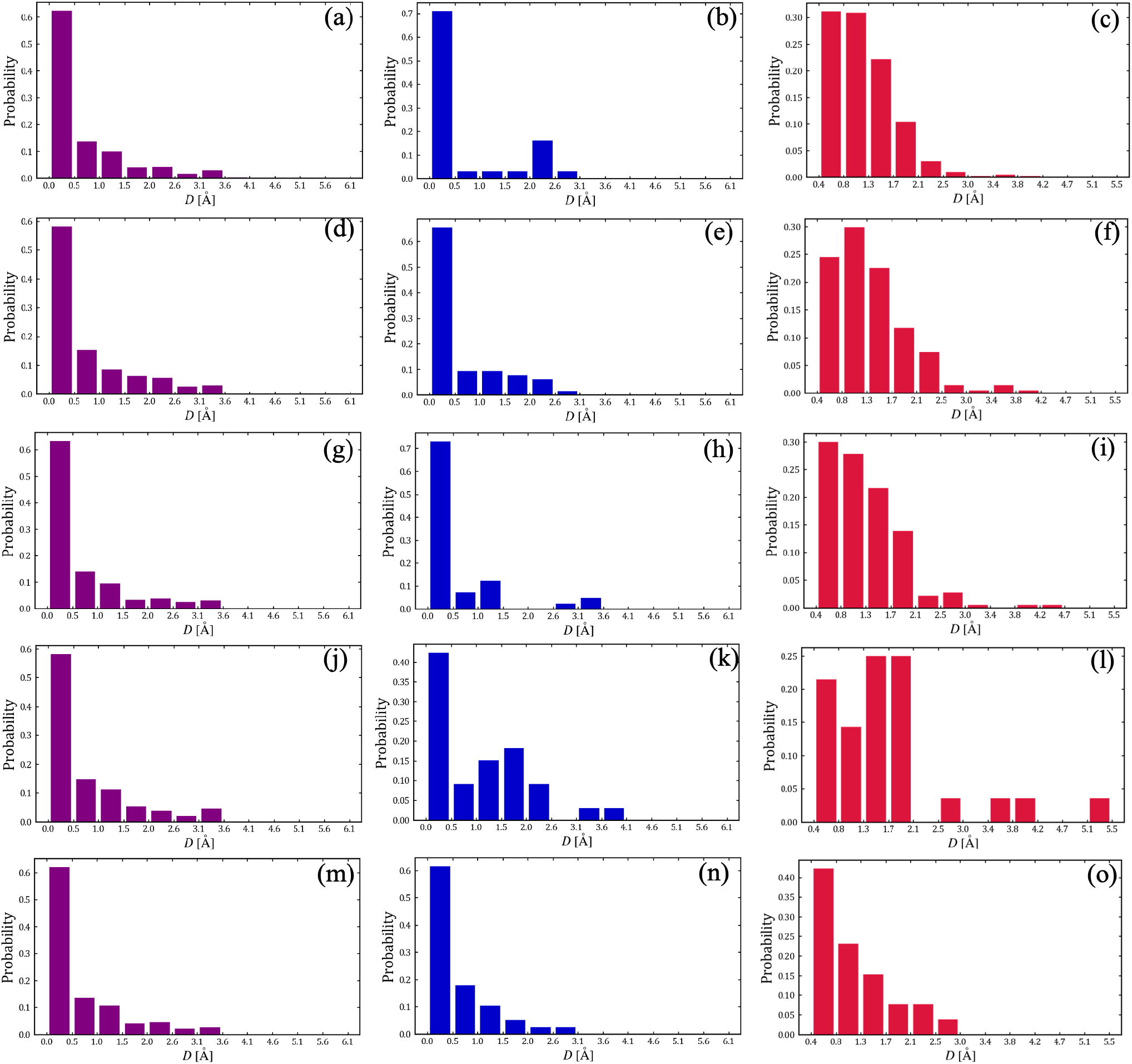
Histograms of the *D*_*i*_ values (Eq. (5)) for five proteins, namely (a), (b), and (c) 2ha2; (d), (e), and (f) 2o4l; (g), (h), and (i) 3jvr; (j), (k), and (l) 3o5n; (m), (n), and (o) 4kao. For each protein, *D*_*i*_ was computed for the oxygen atoms at the positions with *g*_O_(***r***) > 1.5 [(a), (d), (g), (j), and (m)] and those in the ligand-binding pocket [(b), (e), (h), (k), and (n)], and for the crystal waters [(c), (f), (i), (l), and (o)].

**Table 4.** Results using the program Placevent for the five proteins.

| PDB<br>code | Placevent<br>$g_o(\mathbf{r}) > 1.5$ | | | Placevent<br>Ligand-binding pocket | | | Placevent<br>Crystal waters | |
| --- | --- | --- | --- | --- | --- | --- | --- | --- |
| | Average and<br>SD of $D_i$ (Å) | $N_{\text{RISM}}$ | $N_{\text{Model}}$ | Average and<br>SD of $D_i$ (Å) | $N_{\text{RISM}}$ | $N_{\text{Model}}$ | Average and<br>SD of $D_i$ (Å) | $N_{\text{CW}}$ |
| 2ha2 | 0.625±0.888 | 2326 | 2204 | 0.601±0.968 | 31 | 29 | 1.089±0.331 | 436 |
| 2o4l | 0.704±0.924 | 753 | 701 | 0.558±0.746 | 64 | 62 | 1.259±0.452 | 217 |
| 3jvr | 0.607±0.876 | 1530 | 1458 | 0.509±0.839 | 41 | 41 | 1.141±0.451 | 203 |
| 3o5n | 0.705±0.935 | 682 | 630 | 1.066±1.000 | 33 | 35 | 1.674±1.206 | 28 |
| 4kao | 0.626±0.860 | 1480 | 1400 | 0.542±0.671 | 39 | 37 | 0.994±0.427 | 30 |

**Fig. 7.**
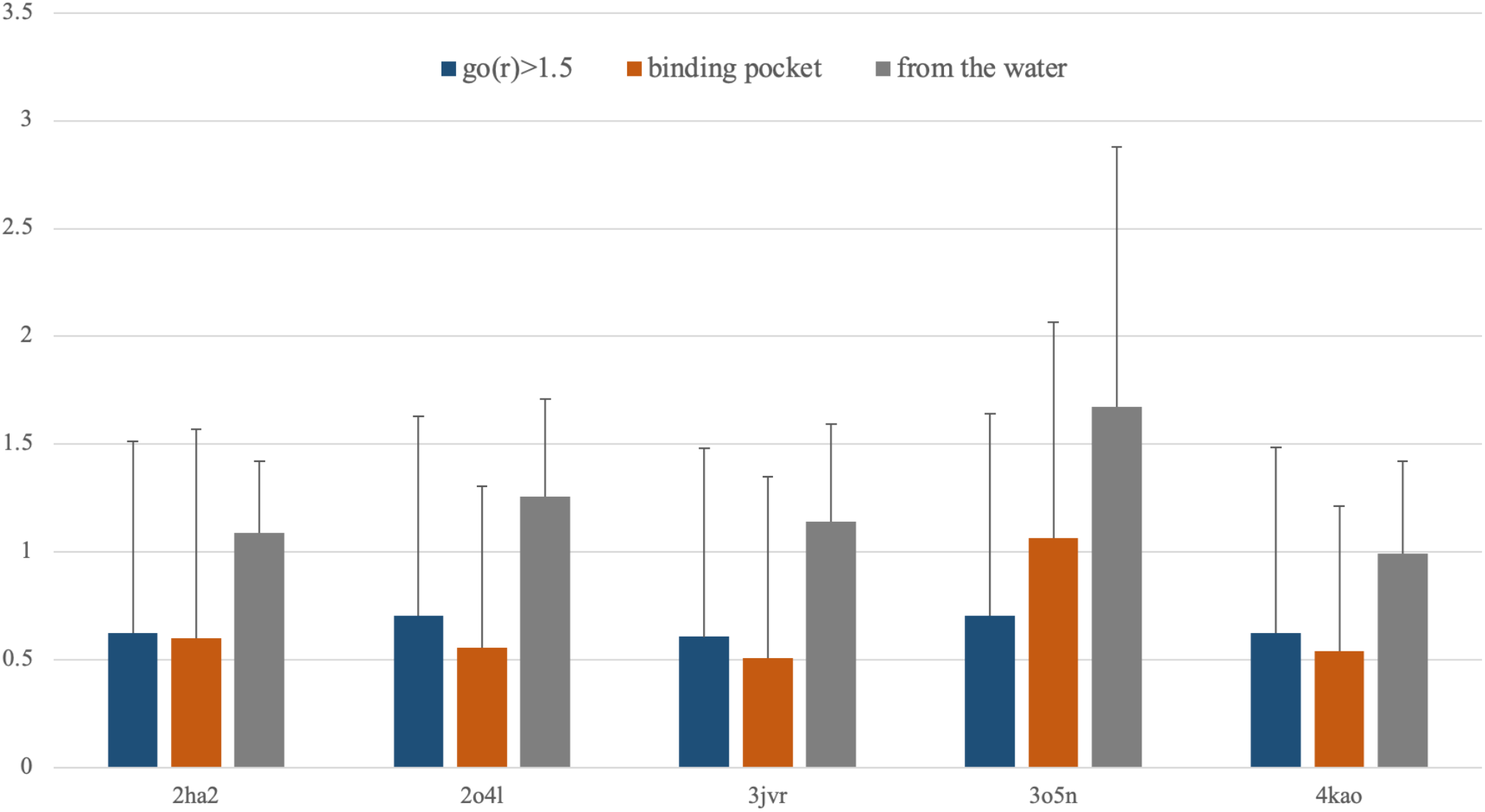
Average and standard deviation of the *D*_*i*_ values for the five proteins.

Nevertheless, the *N*_Model_ values were different from the *N*_RISM_ values (Table 4), reasonably because the peak height and peak position of *g*_O_(***r***) were slightly different in the two methods. Particularly, the *N*_Model_ values were smaller than the corresponding *N*_RISM_ value for all the proteins because our DL model predicted smaller peak values of *g*_O_(***r***) than those obtained using the 3D-RISM theory.

The water placement results of our DL model were afterwards compared with the positions of the crystal waters. To this end, *D*_*i*_ was computed for the crystal waters within 5 Å from the heavy atoms of the protein. As shown in Table 4 and Fig. 6 and 7, the average of *D*_*i*_ (1.0–1.6 Å for all five proteins) was 1/2–1/3 of the Lennard-Jones sigma value of the water oxygen atom for the coincident SPC/E model, indicating that the positions of water oxygen atoms obtained using our DL model were close to those of crystal waters.

### Prediction of the distribution function of water hydrogen sites

A DL model for predicting *g*_H_(***r***) was constructed using the same U-net architecture as that used in the deep-learning model for *g*_O_(***r***). The optimized hyperparameter set (44 in Table S1) was selected considering that the prediction results were not sensitive to this factor (text S3 in Supporting Information). After training the DL model for *g*_H_(***r***) using the optimized hyperparameter set and the twenty-two proteins described in Table 1, the DL model for *g*_H_(***r***) was applied to the five proteins described in Table 1 to predict *g*_H_(***r***). The correlation between the predicted *g*_H_(***r***) values and the *g*_H_(***r***) values obtained using the 3D-RISM theory is reported in Fig. 8, together with the R^2^ score and RMSE values. The DL model for predicting *g*_H_(***r***) exhibited an analogous performance as that for predicting *g*_O_(***r***). Additionally, the RMSE values of *g*_H_(***r***) were smaller than those of *g*_O_(***r***).

**Fig. 8.**
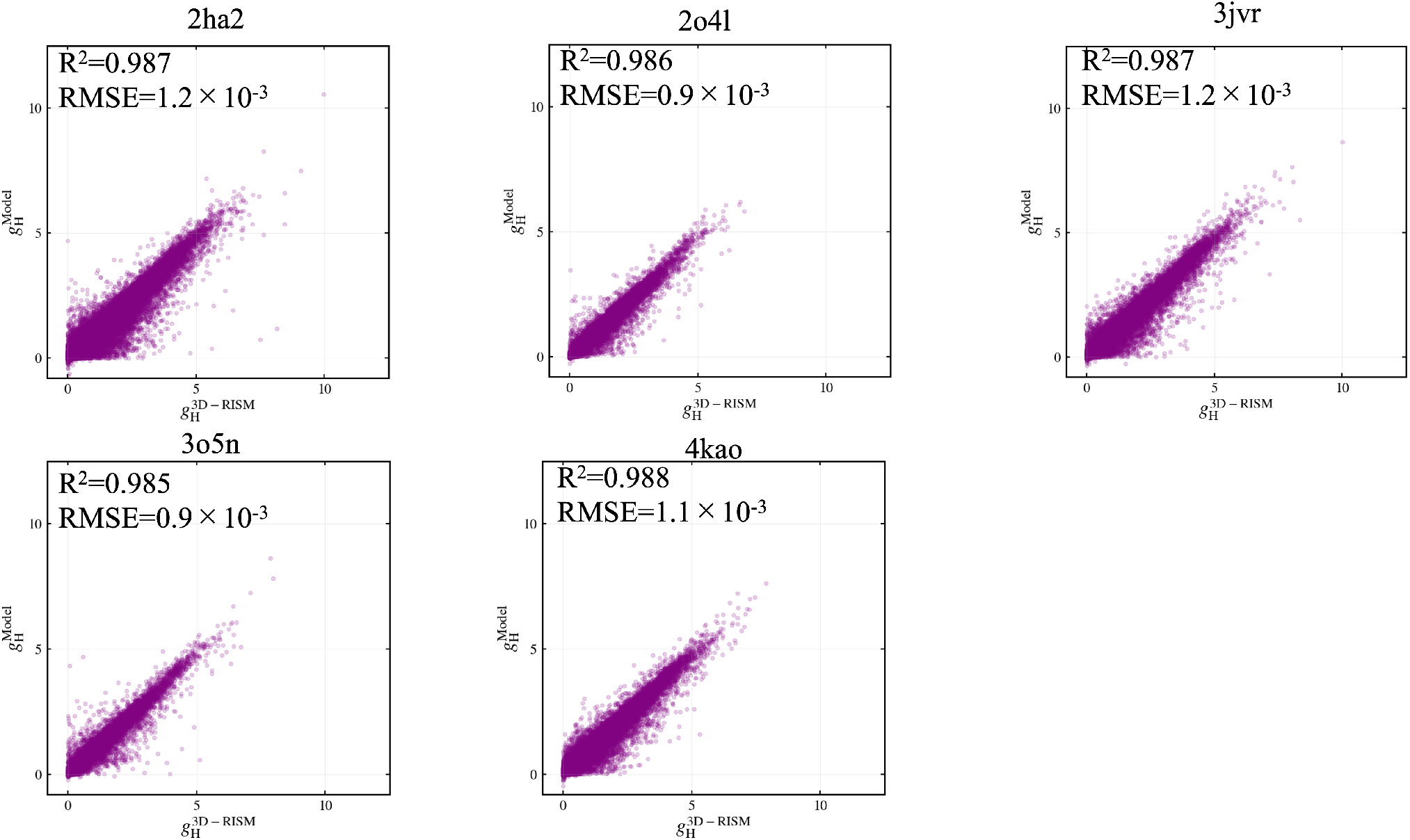
Comparison of the *g*_H_(***r***) values obtained using the 3D-RISM theory and those obtained with our deep-learning model for (a) mouse acetylcholinesterase (PDB code: 2ha2), (b) HIV-1 Protease (PDB code: 2o4l), (c) Checkpoint kinase 1 (PDB code: 3jvr), (d) shank3 PDZ domain (PDB code: 3o5n), and (e) focal adhesion kinase (PDB code: 4kao). The coefficient of determination R^2^-score values and root mean square error (RMSE) are indicated.

### Selection of twenty-seven proteins

To investigate the possible effects of the selection of the proteins on the performance of our DL model, two analyses were conducted.

First, our DL model was applied to the prediction of *g*_O_(***r***) for the 2,691 proteins that were not involved in the twenty-seven proteins in Table 1. The PDB codes of the 2,691 proteins, their R^2^ score values, and their classes are summarized in “Data2718-SI-Forsubmit.xlsx”. The R^2^ score values for the 2691 proteins and the five test proteins were larger than 0.98 for all proteins (Fig. 9). Therefore, our DL model can successfully be applied to various proteins.

**Fig. 9.**
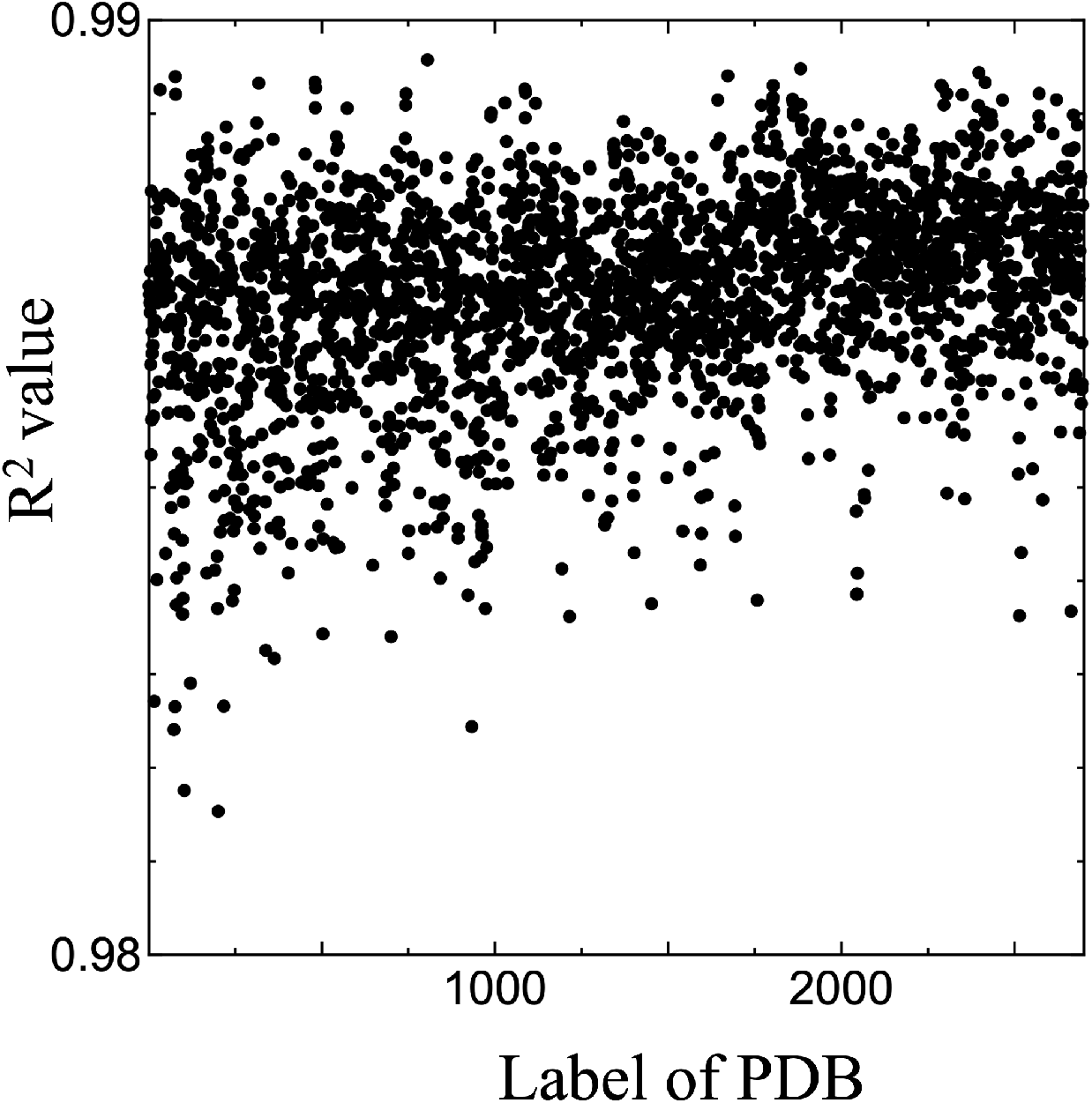
R^2^-score values for the 2696 proteins. The label of each PDB is reported in the file “Data2718-SI-Forsubmit.xlsx”.

In the second analysis, a different pool of twenty-seven proteins was randomly selected for the training and test (Table S4). Twenty-two proteins were used for the training of the DL model for predicting *g*_O_(***r***), and the remaining five proteins were used for the test, with set 44 in Table 3 adopted as hyperparameter set. As shown in Fig. S3, the prediction performance was comparable to that shown in Fig. 3, indicating that the negligible effects of the selection of the twenty-seven proteins on the performance of our DL model.

Therefore, the selection of the twenty-seven proteins shown in Table 1 did not affect the performance of our deep-learning model.

### Comparison of our deep-learning model with other related methodologies

Our DL model was compared with three related methodologies. Two of the three are the method for obtaining the hydration structures around proteins within a short computation time^23,24^. The other is the hybrid method of a DL and the 3D-RISM theory^25^.

First, our DL model is compared with the hybrid method of a DL and the 3D-RISM theory proposed by Sosnin *et al*^25^. Contrarily to our DL model directly predicting *g*_O_(***r***) and *g*_H_(***r***), Sosnin *et al*. proposed a DL model for predicting the bioconcentration-factor values of organic molecules with the input of *g*_O_(***r***) and *g*_H_(***r***) obtained with the 3D-RISM theory. The employed DL model was also different: Sosnin *et al*. employed a three-dimensional convolutional neural network.

Ghanbarpour *et al*.^23^ proposed a DL model for predicting the hydration structure around the proteins. In their study, the hydration structure was characterized by the water occupancy, namely the probability that a water molecule is found at a given grid position. From the definition of the water occupancy, it is closely related to *g*_O_(***r***). Although Ghanbarpour *et al*. attempted to predict the water occupancies using the model based on the U-net architecture, the prediction performance was unsatisfactory. Therefore, they proposed another regression model to predict the water occupancies. However, their model required a preliminary classification using the model predicting the grid points into those high and low water occupancies. Such classification was not required in our DL model.

Maruyama and Hirata^24^ have proposed a fast algorithm to accelerate the 3D-RISM calculation using GPU. The computation of the 3D-RISM calculation for a single protein was finished within a few minutes with a Tesla-K40 GPU^26^. Compared with the algorithm proposed by Maruyama and Hirata, our DL model had two advantages. First, even with a single CPU, the computation was rapidly completed (a few minutes). Furthermore, our DL model enabled to compute *g*_O_(***r***) at a focused region in the protein, such as the ligand-binding pocket or another region of interest, because the protein was decomposed into small boxes of 48^3^ voxels. Such computation is unfeasible for the 3D-RISM theory.

## 4. CONCLUSIONS

In the present study, we proposed a DL model for predicting the hydration structure around the protein based on the U-net architecture. The output was the distribution function of water oxygen *g*_O_(***r***) and hydrogen *g*_H_(***r***)solely with the input of the protein 3D structure.

Our DL model successfully reproduced *g*_O_(***r***) and *g*_H_(***r***)obtained using the 3D-RISM theory of five proteins not included in the training set. The coefficient of determination, R^2^-score values were approximately 0.98 for the five proteins, indicating the good performance of our DL model. Moreover, the model accurately predicted the peak positions of *g*_O_(***r***) from the comparison of the positions of the water oxygen atoms, using Placevent, between our DL model and the 3D-RISM theory. The average of *D*_*i*_ (0.6–0.7 Å), which is the distance of water molecules between that placed by the 3D-RISM theory and the one predicted by our DL model, was small compared to the size of the water oxygen atom, 3 Å. Our DL model also successfully predicted *g*_H_(***r***). In summary, our DL model exhibited a good prediction performance for *g*_O_(***r***) and *g*_H_(***r***).

For the whole protein, our DL model predicted *g*_O_(***r***) within a minute using a single GPU on average. Moreover, *g*_O_(***r***) was predicted for only a focused region of interest, such as the ligand binding domain.

One of the limitations of our DL model is the restricted atom types that can be included, namely carbon, nitrogen, oxygen, sulfur, and hydrogen. Therefore, the application of the current DL model to protein systems involving other atoms (e.g., metals, phosphorus of phosphorylated amino acids, selenium of selenomethione, ions, halogens of ligands, and co-factors) is unfeasible. To extend the applicability of our DL model, the number of atom types should be increased. The data including these atom types and training of our DL model are the object of our future publication.

## DATA AND SOFTWARE AVAILABILITY

Our program, named “gr Predictor”, is available under the GNU General Public License from https://github.com/YoshidomeGroup-Hydration/gr-predictor. Usage of the program is described in the web page described above. All the data used in the present study have been exhaustively presented in the manuscript.

## ASSOCIATED CONTENT

### Supporting Information

The following files are available free of charge.

brief description (file type, i.e., PDF)

## AUTHOR INFORMATION

### Authors

**Kousuke Kawama** - Department of Applied Physics, Graduate School of Engineering, Tohoku University, Sendai 980-8579, Japan.

**Yusaku Fukushima** - Department of Applied Physics, Graduate School of Engineering, Tohoku University, Sendai 980-8579, Japan.

**Mitsunori Ikeguchi** - Graduate School of Medical Life Science, Yokohama City University, 1-7-29, Suehiro-cho, Tsurumi-ku, Yokohama 230-0045, Japan, and AI-driven Drug Discovery Collaborative Unit, HPC- and AI-driven Drug Development Platform Division Center for Computational Science, RIKEN, 1-7-22 Suehiro-cho, Tsurumi-ku, Yokohama, Kanagawa, 230-0045, Japan;.

**Masateru Ohta** - AI-driven Drug Development Platform Division Center for Computational Science, RIKEN, 1-7-22 Suehiro-cho, Tsurumi-ku, Yokohama, Kanagawa, 230-0045, Japan;.

## Author Contribution

T.Y., M.O., and M.I. designed the study. K.K. and Y.F. performed the computations, and K.K., Y.F., and T.Y. analyzed the data. T.Y. and M.O. wrote the article.

## ACKNOWLEDGMENT

Part of the computation was carried out using the computer resource offered under the category of HPCI System Research Project (Project ID: hp210081) by Research Institute for Information Technology, Kyushu University. T.Y. thanks to Mr. Dan Ohashi for the computation of the prediction of *g*_O_(***r***) for 2,696 proteins.

