## Supporting Information for "gr Predictor: a Deep-Learning Model for Predicting the Hydration Structures around Proteins"

**SI text S1. Similarities between the sequences of the proteins used in the study**

To discuss the similarities between the sequences of the proteins used in the present study, the clustering of the protein sequences in accordance with the sequence similarity was performed using the program Cd-hit [S1] on 2,718 proteins. The sequence identity threshold was set at 0.9, indicating that the proteins with sequence similarity larger than 90% was classified in the same cluster.

The 2,718 proteins were classified into 863 classes, summarized in the file "Data2718-SI-Forsubmit.xlsx".

The twenty-seven proteins of this study were classified into twenty-five clusters. The following PDB pairs were classified in the same clusters and used for the training: 3drf and 3dri; and 4ezr and 4ezz. None of the proteins used in the test had 90% sequence similarity to the twenty-two proteins used for in the training.

[S1] Li W., Adam Godzik A., Cd-hit: a fast program for clustering and comparing large sets of protein or nucleotide sequences, *Bioinformatics* **2006**, *22*, 1658-1659.

**SI text S2. Effect of the convolution in the upsampling layers on the performance of the deep-learning model**

The convolution followed by the upsampling in the original U-net architecture was removed in our architecture. To discuss the effect of the convolution on the performance of our deep-learning model, the model in which the convolution was taken into consideration was also constructed. The deep-learning model for predicting $g_0(\boldsymbol{r})$ was constructed. The optimized hyperparameter set was number 44 in Table S2. The $R^2$ values for the five proteins are shown in Table S3. The values were comparable to those obtained using our DL model. Therefore, the effect of the convolution on the performance of our deep-learning model was negligible.

### SI text S3. Results of the cross validation

The $\bar{E}_{\mathrm{Validation}}(200)$ value in Table S2 was less than 0.01 for 31% of the hyperparameter sets. From the definition of the loss function in Eq. (3) in the main text, the average of the deviation of $g_{\mathrm{O}}(\boldsymbol{r})$ obtained by our DL model from $g_{\mathrm{O}}(\boldsymbol{r})$ obtained by the 3D-RISM theory was 0.1. For these hyperparameter sets, $E_{\mathrm{Training}}$ and $E_{\mathrm{Validation}}$ converged during the 200-epochs training (Fig. S2). For the other hyperparameter sets, the $\bar{E}_{\mathrm{Validation}}(200)$ value was larger than 0.01 because $E_{\mathrm{Training}}$ did not converge during 200-epochs training. However, for these hyperparameter sets, we did not perform training with larger epochs than 200, as the $\bar{E}_{\mathrm{Validation}}(200)$ value would be expected to converge to ~0.01.

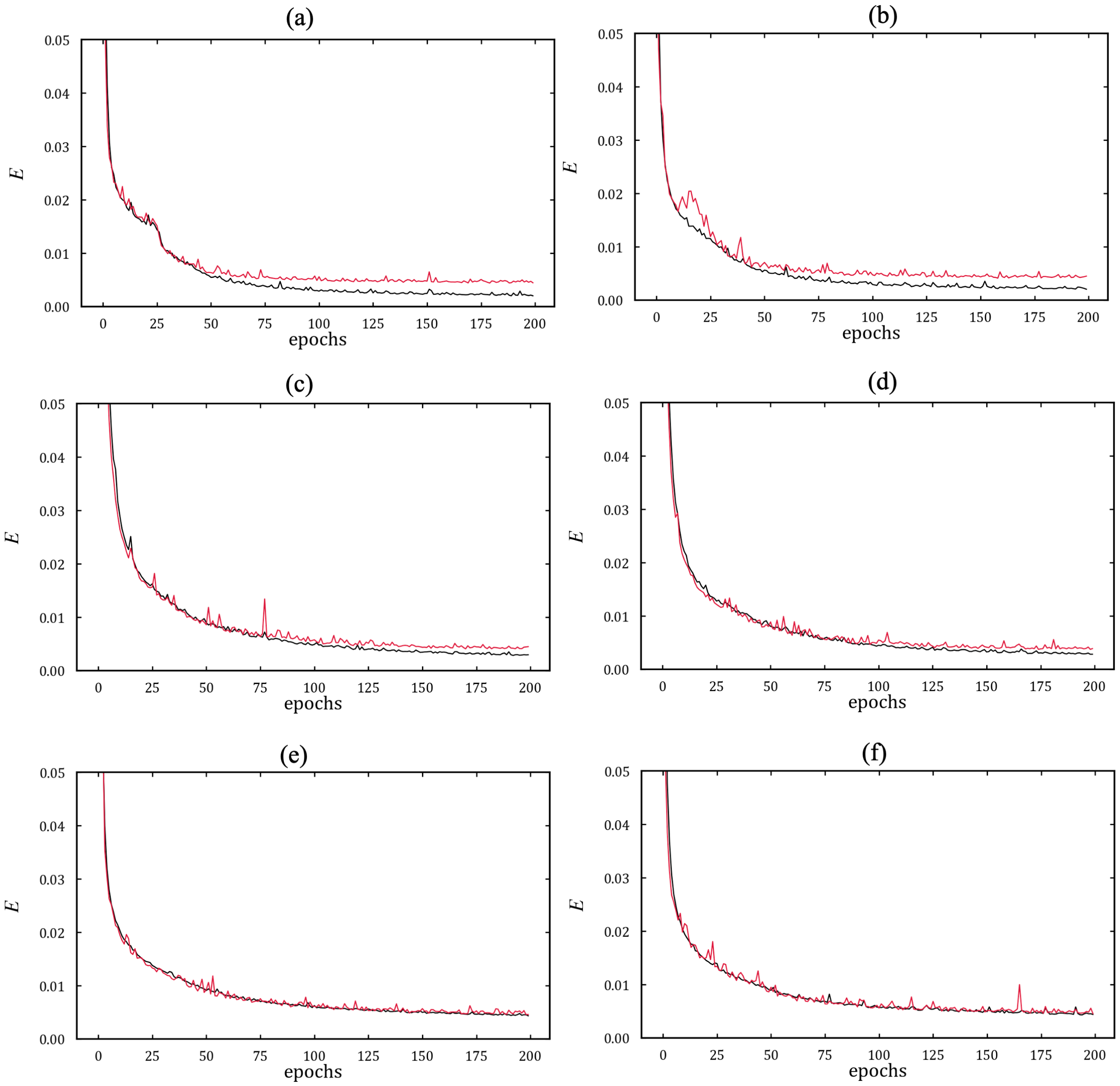


**Fig. S1.** Correlation of the loss function, $E$, defined by Eq. (3) in the main text, to the number of epochs. In each subpart, the loss function for the training and validation are indicated with black and red lines, respectively. For the computation of $E$ in the validation, the $E$ value at $i$ epoch was obtained using the model at $i$ epoch. In (a), (c), and (e), the ten proteins and twelve proteins described in Table 1 were used for the training and validation, respectively. In (b), (d), and (f), the twelve proteins and the ten proteins were used for the training and validation, respectively. The hyperparameter set numbers shown in Table S2 are 46 [(a) and (b)], 44 [(c) and (d)], and 17 [(e) and (f)].

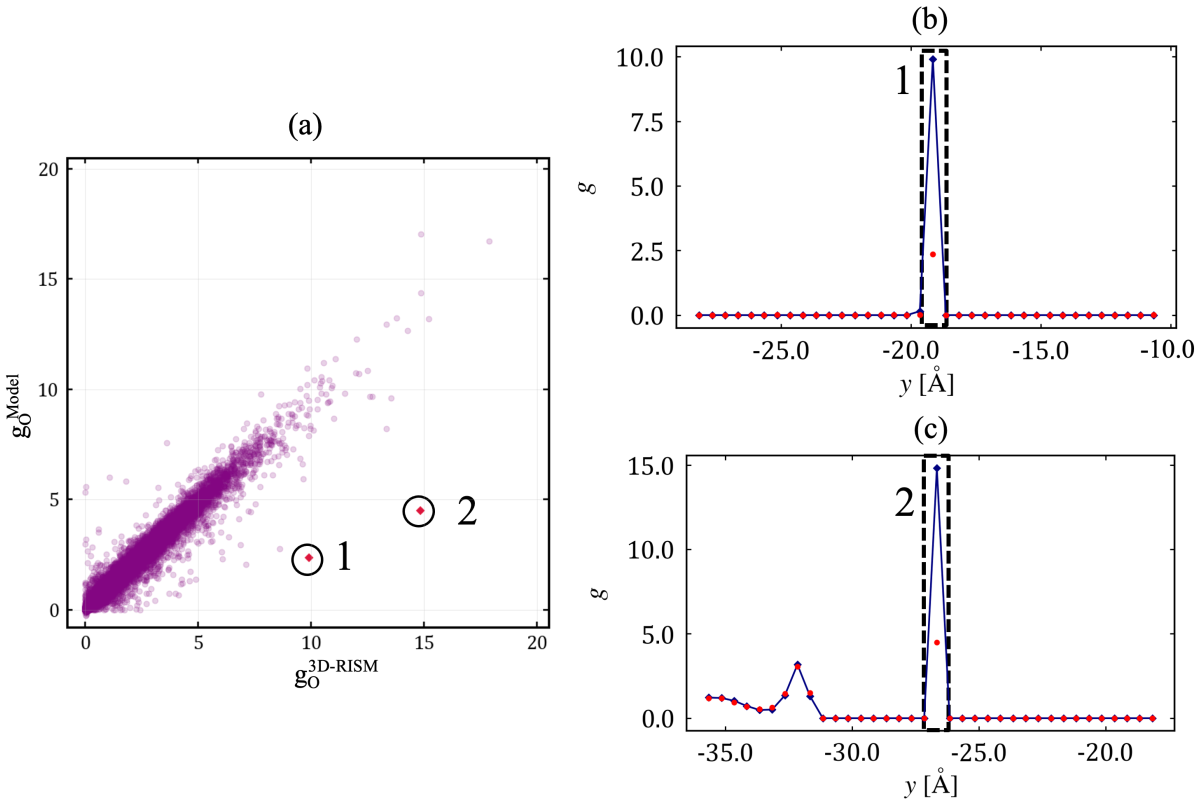


**Fig. S2.** Examples illustrating that the $g_O(\boldsymbol{r})$ values obtained using our deep-learning model were substantially different from those obtained with the 3D-RISM theory. The protein was shank3 PDZ domain (PDB code: 3o5n). (a) Same figure as Fig. 3 in the main text. The $g_O(\boldsymbol{r})$ values at the points labeled by "1" and "2" are shown in (b) and (c). The axis is defined in Fig. 5 in the main text. In the figure, the blue points and lines represent $g_O^{3D-RISM}(\boldsymbol{r})$, whereas the red points represent $g_O^{Model}(\boldsymbol{r})$. Because the $g_O(\boldsymbol{r})$ values at most of the points in (b) and (c) are zero, the regions plotted in (b) and (c) correspond to the interior of the protein.

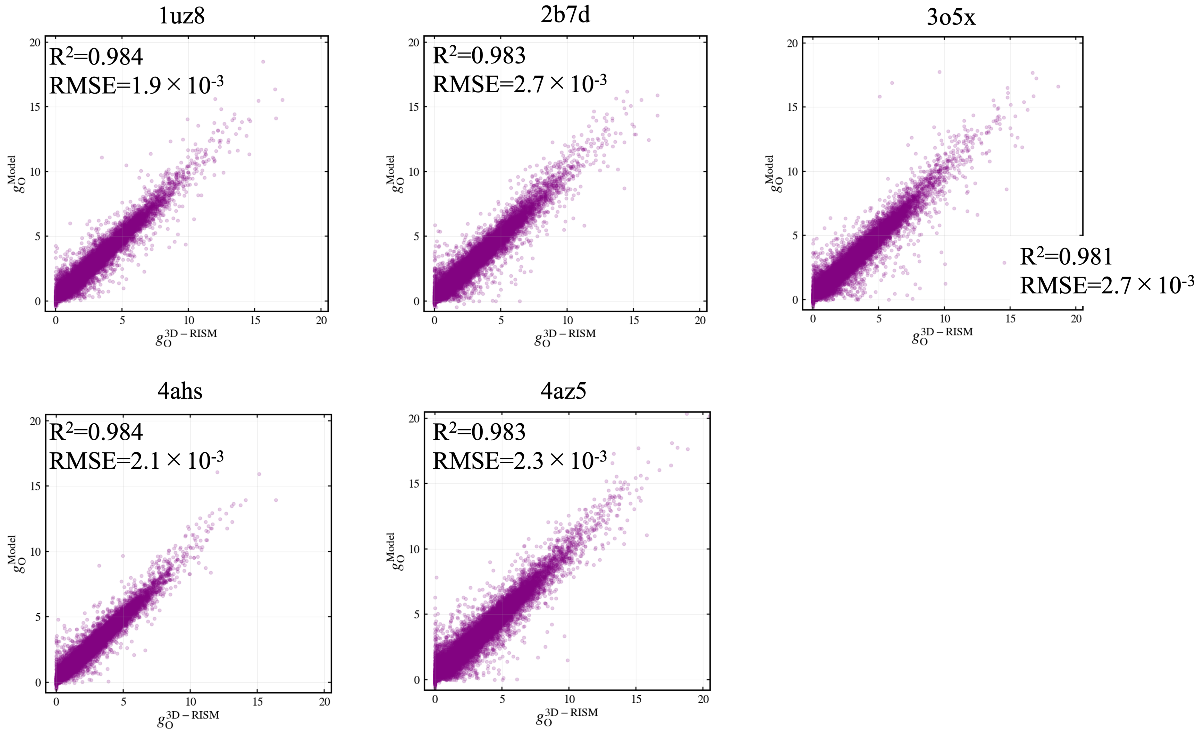


**Fig. S3.** Correlation between the $g_O(\boldsymbol{r})$ values predicted by our deep-learning model and the values calculated by the 3D-RISM theory. The twenty-seven proteins used for the training and test are summarized in Table S4. The coefficient of determination $R^2$-score values and root mean square error (RMSE) are indicated.

**Table S1.** $\sigma_{\mathrm{vdw},j}$ value for each atom type.

| | $\sigma_{\mathrm{vdw},j}$ (Å) |
|---|---|
| Carbon | 1.69984 |
| Oxygen | 1.51369 |
| Nitrogen | 1.62500 |
| Sulfur | 1.78180 |
| Hydrogen | 1.20000 |

**Table S2**. Values of the error functions for the 162 hyperparameter sets of the deep-learning model for $g_O(\boldsymbol{r})$. In the table, the $E_{Validation}(200)$ value obtained using the ten proteins is denoted by $E^1_{\mathrm{Valid}}$, and that obtained using the twelve proteins is denoted by $E^2_{\mathrm{Valid}}$. The average of $E^1_{\mathrm{Valid}}$ and $E^2_{\mathrm{Valid}}$ is also shown. "A", "B", "C", and "D" represent the hyperparameter shown in Table 3 in the main text. The definitions of the layers are reported in Fig. 2 in the main text. The hyperparameter set number for the optimized hyperparameter set (in blue) was 44.

| Hyperparameter set number | A | B | C | D | | $E^1_{\mathrm{Valid}}$ | $E^2_{\mathrm{Valid}}$ | Average of $E^1_{\mathrm{Valid}}$ and $E^2_{\mathrm{Valid}}$ |
|---|---|---|---|---|---|---|---|---|
| 0 | 3 | 16 | - | None | None | 0.0070 | 0.0054 | 0.0062 |
| 1 | 3 | 16 | 0.3 | - | 5th layer | 0.0096 | 0.0068 | 0.0082 |
| 2 | 3 | 16 | 0.5 | - | 5th layer | 0.0075 | 0.0067 | 0.0071 |
| 3 | 3 | 16 | 0.3 | i | 4th-5th layers | 0.0056 | 0.0070 | 0.0063 |
| 4 | 3 | 16 | 0.3 | ii | 4th-5th layers | 0.0068 | 0.0079 | 0.0074 |
| 5 | 3 | 16 | 0.3 | iii | 4th-5th layers | 0.0075 | 0.0075 | 0.0075 |
| 6 | 3 | 16 | 0.5 | i | 4th-5th layers | 0.0063 | 0.0085 | 0.0074 |
| 7 | 3 | 16 | 0.5 | ii | 4th-5th layers | 0.0068 | 0.0069 | 0.0068 |
| 8 | 3 | 16 | 0.5 | iii | 4th-5th layers | 0.0066 | 0.0068 | 0.0067 |
| 9 | 3 | 16 | 0.3 | i | 3rd-5th layers | 0.0057 | 0.0066 | 0.0061 |
| 10 | 3 | 16 | 0.3 | ii | 3rd-5th layers | 0.0062 | 0.0055 | 0.0059 |
| 11 | 3 | 16 | 0.3 | iii | 3rd-5th layers | 0.0054 | 0.0054 | 0.0054 |
| 12 | 3 | 16 | 0.5 | i | 3rd-5th layers | 0.0064 | 0.0060 | 0.0062 |
| 13 | 3 | 16 | 0.5 | ii | 3rd-5th layers | 0.0064 | 0.0066 | 0.0065 |
| 14 | 3 | 16 | 0.5 | iii | 3rd-5th layers | 0.0065 | 0.0078 | 0.0071 |
| 15 | 3 | 16 | 0.3 | i | 2nd-5th layers | 0.0055 | 0.0048 | 0.0052 |
| 16 | 3 | 16 | 0.3 | ii | 2nd-5th layers | 0.0051 | 0.0048 | 0.0050 |
| 17 | 3 | 16 | 0.3 | iii | 2nd-5th layers | 0.0046 | 0.0046 | 0.0046 |
| 18 | 3 | 16 | 0.5 | i | 2nd-5th layers | 0.0056 | 0.0046 | 0.0051 |
| 19 | 3 | 16 | 0.5 | ii | 2nd-5th layers | 0.0050 | 0.0054 | 0.0052 |
| 20 | 3 | 16 | 0.5 | iii | 2nd-5th layers | 0.0070 | 0.0079 | 0.0074 |
| 21 | 3 | 16 | 0.3 | i | all layers | 0.0060 | 0.0061 | 0.0061 |
| 22 | 3 | 16 | 0.3 | ii | all layers | 0.0082 | 0.0080 | 0.0081 |
| 23 | 3 | 16 | 0.3 | iii | all layers | 0.0094 | 0.0124 | 0.0109 |
| 24 | 3 | 16 | 0.5 | i | all layers | 0.0105 | 0.0105 | 0.0105 |
| 25 | 3 | 16 | 0.5 | ii | all layers | 0.0117 | 0.0110 | 0.0114 |
| 26 | 3 | 16 | 0.5 | iii | all layers | 0.0272 | 0.0248 | 0.0260 |

| | | | | | | | | |
|---|---|---|---|---|---|---|---|---|
| 27 | 3 | 32 | - | None | None | 0.0209 | 0.0148 | 0.0178 |
| 28 | 3 | 32 | 0.3 | - | $5^{th}$ layer | 0.0151 | 0.0101 | 0.0126 |
| 29 | 3 | 32 | 0.5 | - | $5^{th}$ layer | 0.0072 | 0.0094 | 0.0083 |
| 30 | 3 | 32 | 0.3 | i | $4^{th}$-$5^{th}$ layers | 0.0078 | 0.0085 | 0.0082 |
| 31 | 3 | 32 | 0.3 | ii | $4^{th}$-$5^{th}$ layers | 0.0061 | 0.0136 | 0.0098 |
| 32 | 3 | 32 | 0.3 | iii | $4^{th}$-$5^{th}$ layers | 0.0072 | 0.0070 | 0.0071 |
| 33 | 3 | 32 | 0.5 | i | $4^{th}$-$5^{th}$ layers | 0.0075 | 0.0079 | 0.0077 |
| 34 | 3 | 32 | 0.5 | ii | $4^{th}$-$5^{th}$ layers | 0.0072 | 0.0078 | 0.0075 |
| 35 | 3 | 32 | 0.5 | iii | $4^{th}$-$5^{th}$ layers | 0.0047 | 0.0048 | 0.0047 |
| 36 | 3 | 32 | 0.3 | i | $3^{rd}$-$5^{th}$ layers | 0.0074 | 0.0075 | 0.0074 |
| 37 | 3 | 32 | 0.3 | ii | $3^{rd}$-$5^{th}$ layers | 0.0111 | 0.0097 | 0.0104 |
| 38 | 3 | 32 | 0.3 | iii | $3^{rd}$-$5^{th}$ layers | 0.0053 | 0.2409 | 0.1231 |
| 39 | 3 | 32 | 0.5 | i | $3^{rd}$-$5^{th}$ layers | 0.0110 | 0.0064 | 0.0087 |
| 40 | 3 | 32 | 0.5 | ii | $3^{rd}$-$5^{th}$ layers | 0.0181 | 0.0072 | 0.0127 |
| 41 | 3 | 32 | 0.5 | iii | $3^{rd}$-$5^{th}$ layers | 0.0049 | 0.0045 | 0.0047 |
| 42 | 3 | 32 | 0.3 | i | $2^{nd}$-$5^{th}$ layers | 0.0044 | 0.0046 | 0.0045 |
| 43 | 3 | 32 | 0.3 | ii | $2^{nd}$-$5^{th}$ layers | 0.0049 | 0.0048 | 0.0048 |
| **44** | **3** | **32** | **0.3** | **iii** | $2^{nd}$-$5^{th}$ layers | **0.0045** | **0.0039** | **0.0042** |
| 45 | 3 | 32 | 0.5 | i | $2^{nd}$-$5^{th}$ layers | 0.0091 | 0.0065 | 0.0078 |
| 46 | 3 | 32 | 0.5 | ii | $2^{nd}$-$5^{th}$ layers | 0.0045 | 0.0045 | 0.0045 |
| 47 | 3 | 32 | 0.5 | iii | $2^{nd}$-$5^{th}$ layers | 0.0617 | 0.0065 | 0.0341 |
| 48 | 3 | 32 | 0.3 | i | all layers | 0.0106 | 0.0142 | 0.0124 |
| 49 | 3 | 32 | 0.3 | ii | all layers | 0.0059 | 0.0063 | 0.0061 |
| 50 | 3 | 32 | 0.3 | iii | all layers | 0.0074 | 0.0063 | 0.0069 |
| 51 | 3 | 32 | 0.5 | i | all layers | 0.0170 | 0.0240 | 0.0205 |
| 52 | 3 | 32 | 0.5 | ii | all layers | 0.0133 | 0.0097 | 0.0115 |
| 53 | 3 | 32 | 0.5 | iii | all layers | 0.0161 | 0.0174 | 0.0168 |
| 54 | 4 | 16 | - | None | None | 0.0191 | 0.0174 | 0.0182 |
| 55 | 4 | 16 | 0.3 | - | $5^{th}$ layer | 0.0175 | 0.0173 | 0.0174 |
| 56 | 4 | 16 | 0.5 | - | $5^{th}$ layer | 0.0319 | 0.0181 | 0.0250 |
| 57 | 4 | 16 | 0.3 | i | $4^{th}$-$5^{th}$ layers | 0.0237 | 0.0229 | 0.0233 |
| 58 | 4 | 16 | 0.3 | ii | $4^{th}$-$5^{th}$ layers | 0.0201 | 0.0177 | 0.0189 |
| 59 | 4 | 16 | 0.3 | iii | $4^{th}$-$5^{th}$ layers | 0.0301 | 0.0316 | 0.0309 |
| 60 | 4 | 16 | 0.5 | i | $4^{th}$-$5^{th}$ layers | 0.0232 | 0.0285 | 0.0259 |
| 61 | 4 | 16 | 0.5 | ii | $4^{th}$-$5^{th}$ layers | 0.0260 | 0.0197 | 0.0229 |
| 62 | 4 | 16 | 0.5 | iii | $4^{th}$-$5^{th}$ layers | 0.0516 | 0.0767 | 0.0641 |
| 63 | 4 | 16 | 0.3 | i | $3^{rd}$-$5^{th}$ layers | 0.0949 | 0.0318 | 0.0634 |
| 64 | 4 | 16 | 0.3 | ii | $3^{rd}$-$5^{th}$ layers | 0.0261 | 0.0749 | 0.0505 |

| | | | | | | | | |
|---|---|---|---|---|---|---|---|---|
| 65 | 4 | 16 | 0.3 | iii | 3rd-5th layers | 0.0768 | 0.0236 | 0.0502 |
| 66 | 4 | 16 | 0.5 | i | 3rd-5th layers | 0.0165 | 0.0694 | 0.0430 |
| 67 | 4 | 16 | 0.5 | ii | 3rd-5th layers | 0.0369 | 0.0478 | 0.0423 |
| 68 | 4 | 16 | 0.5 | iii | 3rd-5th layers | 0.0066 | 0.0074 | 0.0070 |
| 69 | 4 | 16 | 0.3 | i | 2nd-5th layers | 0.0103 | 0.0261 | 0.0182 |
| 70 | 4 | 16 | 0.3 | ii | 2nd-5th layers | 0.0109 | 0.0140 | 0.0125 |
| 71 | 4 | 16 | 0.3 | iii | 2nd-5th layers | 0.0099 | 0.0059 | 0.0079 |
| 72 | 4 | 16 | 0.5 | i | 2nd-5th layers | 0.0163 | 0.0070 | 0.0116 |
| 73 | 4 | 16 | 0.5 | ii | 2nd-5th layers | 0.0100 | 0.0139 | 0.0119 |
| 74 | 4 | 16 | 0.5 | iii | 2nd-5th layers | 0.0112 | 0.0119 | 0.0115 |
| 75 | 4 | 16 | 0.3 | i | all layers | 0.0081 | 0.0095 | 0.0088 |
| 76 | 4 | 16 | 0.3 | ii | all layers | 0.0097 | 0.0115 | 0.0106 |
| 77 | 4 | 16 | 0.3 | iii | all layers | 0.0095 | 0.0096 | 0.0096 |
| 78 | 4 | 16 | 0.5 | i | all layers | 0.0202 | 0.0185 | 0.0193 |
| 79 | 4 | 16 | 0.5 | ii | all layers | 0.0160 | 0.0130 | 0.0145 |
| 80 | 4 | 16 | 0.5 | iii | all layers | 0.0229 | 0.0213 | 0.0221 |
| 81 | 4 | 32 | - | None | None | 0.0372 | 0.0785 | 0.0578 |
| 82 | 4 | 32 | 0.3 | - | 5th layer | 0.0927 | 0.0353 | 0.0640 |
| 83 | 4 | 32 | 0.5 | - | 5th layer | 0.0740 | 0.0895 | 0.0818 |
| 84 | 4 | 32 | 0.3 | i | 4th-5th layers | 0.0656 | 0.0778 | 0.0717 |
| 85 | 4 | 32 | 0.3 | ii | 4th-5th layers | 0.0798 | 0.0422 | 0.0610 |
| 86 | 4 | 32 | 0.3 | iii | 4th-5th layers | 0.0665 | 0.0769 | 0.0717 |
| 87 | 4 | 32 | 0.5 | i | 4th-5th layers | 0.0899 | 0.0230 | 0.0565 |
| 88 | 4 | 32 | 0.5 | ii | 4th-5th layers | 0.0835 | 0.0497 | 0.0666 |
| 89 | 4 | 32 | 0.5 | iii | 4th-5th layers | 0.0135 | 0.2409 | 0.1272 |
| 90 | 4 | 32 | 0.3 | i | 3rd-5th layers | 0.0274 | 0.0651 | 0.0463 |
| 91 | 4 | 32 | 0.3 | ii | 3rd-5th layers | 0.0798 | 0.0118 | 0.0458 |
| 92 | 4 | 32 | 0.3 | iii | 3rd-5th layers | 0.0568 | 0.0679 | 0.0624 |
| 93 | 4 | 32 | 0.5 | i | 3rd-5th layers | 0.0112 | 0.0092 | 0.0102 |
| 94 | 4 | 32 | 0.5 | ii | 3rd-5th layers | 0.0133 | 0.0798 | 0.0466 |
| 95 | 4 | 32 | 0.5 | iii | 3rd-5th layers | Failed to converge | 0.0091 | Failed to converge |
| 96 | 4 | 32 | 0.3 | i | 2nd-5th layers | 0.0695 | 0.2409 | 0.1552 |
| 97 | 4 | 32 | 0.3 | ii | 2nd-5th layers | 0.0650 | 0.0123 | 0.0387 |
| 98 | 4 | 32 | 0.3 | iii | 2nd-5th layers | 0.0067 | 0.0070 | 0.0069 |
| 99 | 4 | 32 | 0.5 | i | 2nd-5th layers | 0.0123 | 0.0118 | 0.0121 |
| 100 | 4 | 32 | 0.5 | ii | 2nd-5th layers | 0.0094 | 0.1251 | 0.0673 |
| 101 | 4 | 32 | 0.5 | iii | 2nd-5th layers | 0.2441 | 0.2409 | 0.2425 |

| | | | | | | | | |
|---|---|---|---|---|---|---|---|---|
| 102 | 4 | 32 | 0.3 | i | all layers | 0.2440 | 0.2409 | 0.2425 |
| 103 | 4 | 32 | 0.3 | ii | all layers | 0.0137 | 0.0727 | 0.0432 |
| 104 | 4 | 32 | 0.3 | iii | all layers | 0.2440 | 0.0107 | 0.1273 |
| 105 | 4 | 32 | 0.5 | i | all layers | 0.0364 | 0.0585 | 0.0475 |
| 106 | 4 | 32 | 0.5 | ii | all layers | 0.2440 | 0.0276 | 0.1358 |
| 107 | 4 | 32 | 0.5 | iii | all layers | 0.0138 | 0.1645 | 0.0891 |
| 108 | 5 | 16 | - | None | None | 0.2440 | 0.0139 | 0.1289 |
| 109 | 5 | 16 | 0.3 | - | $5^{th}$ layer | 0.0133 | 0.2409 | 0.1271 |
| 110 | 5 | 16 | 0.5 | - | $5^{th}$ layer | 0.2440 | 0.2409 | 0.2425 |
| 111 | 5 | 16 | 0.3 | i | $4^{th}$-$5^{th}$ layers | 0.2440 | 0.2409 | 0.2425 |
| 112 | 5 | 16 | 0.3 | ii | $4^{th}$-$5^{th}$ layers | 0.2440 | 0.0130 | 0.1285 |
| 113 | 5 | 16 | 0.3 | iii | $4^{th}$-$5^{th}$ layers | 0.2440 | 0.0207 | 0.1324 |
| 114 | 5 | 16 | 0.5 | i | $4^{th}$-$5^{th}$ layers | 0.0166 | 0.2409 | 0.1288 |
| 115 | 5 | 16 | 0.5 | ii | $4^{th}$-$5^{th}$ layers | 0.2440 | 0.2409 | 0.2425 |
| 116 | 5 | 16 | 0.5 | iii | $4^{th}$-$5^{th}$ layers | 0.2440 | 0.2409 | 0.2425 |
| 117 | 5 | 16 | 0.3 | i | $3^{rd}$-$5^{th}$ layers | 0.2440 | 0.2409 | 0.2424 |
| 118 | 5 | 16 | 0.3 | ii | $3^{rd}$-$5^{th}$ layers | 0.0157 | 0.0412 | 0.0284 |
| 119 | 5 | 16 | 0.3 | iii | $3^{rd}$-$5^{th}$ layers | 0.2440 | 0.2409 | 0.2425 |
| 120 | 5 | 16 | 0.5 | i | $3^{rd}$-$5^{th}$ layers | 0.2440 | 0.0162 | 0.1301 |
| 121 | 5 | 16 | 0.5 | ii | $3^{rd}$-$5^{th}$ layers | 0.0097 | 0.0109 | 0.0103 |
| 122 | 5 | 16 | 0.5 | iii | $3^{rd}$-$5^{th}$ layers | 0.0089 | 0.0082 | 0.0086 |
| 123 | 5 | 16 | 0.3 | i | $2^{nd}$-$5^{th}$ layers | 0.2440 | 0.2409 | 0.2425 |
| 124 | 5 | 16 | 0.3 | ii | $2^{nd}$-$5^{th}$ layers | 0.0116 | 0.0094 | 0.0105 |
| 125 | 5 | 16 | 0.3 | iii | $2^{nd}$-$5^{th}$ layers | 0.0114 | 0.2409 | 0.1261 |
| 126 | 5 | 16 | 0.5 | i | $2^{nd}$-$5^{th}$ layers | 0.2440 | 0.2409 | 0.2425 |
| 127 | 5 | 16 | 0.5 | ii | $2^{nd}$-$5^{th}$ layers | 0.0081 | 0.0062 | 0.0072 |
| 128 | 5 | 16 | 0.5 | iii | $2^{nd}$-$5^{th}$ layers | 0.0063 | 0.0070 | 0.0067 |
| 129 | 5 | 16 | 0.3 | i | all layers | 0.2440 | 0.2409 | 0.2425 |
| 130 | 5 | 16 | 0.3 | ii | all layers | 0.2440 | 0.0100 | 0.1270 |
| 131 | 5 | 16 | 0.3 | iii | all layers | 0.0107 | 0.0314 | 0.0210 |
| 132 | 5 | 16 | 0.5 | i | all layers | 0.2440 | 0.2409 | 0.2424 |
| 133 | 5 | 16 | 0.5 | ii | all layers | 0.0127 | 0.0141 | 0.0134 |
| 134 | 5 | 16 | 0.5 | iii | all layers | 0.2440 | 0.0170 | 0.1305 |
| 135 | 5 | 32 | | None | None | 0.1043 | 0.1228 | 0.1136 |
| 136 | 5 | 32 | 0.3 | None | $5^{th}$ layer | 0.1115 | 0.0893 | 0.1004 |
| 137 | 5 | 32 | 0.5 | None | $5^{th}$ layer | 0.1020 | 0.1048 | 0.1034 |
| 138 | 5 | 32 | 0.3 | i | $4^{th}$-$5^{th}$ layers | 0.1232 | 0.0880 | 0.1056 |
| 139 | 5 | 32 | 0.3 | ii | $4^{th}$-$5^{th}$ layers | 0.1103 | 0.1002 | 0.1052 |

| | | | | | | | | |
|---|---|---|---|---|---|---|---|---|
| 140 | 5 | 32 | 0.3 | iii | $4^{th}$-$5^{th}$ layers | 0.0907 | 0.1035 | 0.0971 |
| 141 | 5 | 32 | 0.5 | i | $4^{th}$-$5^{th}$ layers | 0.1001 | 0.1598 | 0.1299 |
| 142 | 5 | 32 | 0.5 | ii | $4^{th}$-$5^{th}$ layers | 0.0919 | 0.1184 | 0.1051 |
| 143 | 5 | 32 | 0.5 | iii | $4^{th}$-$5^{th}$ layers | 0.2027 | 0.0615 | 0.1321 |
| 144 | 5 | 32 | 0.3 | i | $3^{rd}$-$5^{th}$ layers | 0.0901 | 0.0968 | 0.0935 |
| 145 | 5 | 32 | 0.3 | ii | $3^{rd}$-$5^{th}$ layers | 0.1329 | 0.0899 | 0.1114 |
| 146 | 5 | 32 | 0.3 | iii | $3^{rd}$-$5^{th}$ layers | 0.1720 | 0.1159 | 0.1440 |
| 147 | 5 | 32 | 0.5 | i | $3^{rd}$-$5^{th}$ layers | 0.0461 | 0.1148 | 0.0805 |
| 148 | 5 | 32 | 0.5 | ii | $3^{rd}$-$5^{th}$ layers | 0.2351 | 0.1220 | 0.1785 |
| 149 | 5 | 32 | 0.5 | iii | $3^{rd}$-$5^{th}$ layers | 0.0096 | 0.0066 | 0.0081 |
| 150 | 5 | 32 | 0.3 | i | $2^{nd}$-$5^{th}$ layers | 0.0938 | 0.1026 | 0.0982 |
| 151 | 5 | 32 | 0.3 | ii | $2^{nd}$-$5^{th}$ layers | 0.1004 | 0.1499 | 0.1251 |
| 152 | 5 | 32 | 0.3 | iii | $2^{nd}$-$5^{th}$ layers | 0.0124 | 0.0106 | 0.0115 |
| 153 | 5 | 32 | 0.5 | i | $2^{nd}$-$5^{th}$ layers | 0.0857 | 0.0417 | 0.0637 |
| 154 | 5 | 32 | 0.5 | ii | $2^{nd}$-$5^{th}$ layers | 0.0073 | 0.0068 | 0.0070 |
| 155 | 5 | 32 | 0.5 | iii | $2^{nd}$-$5^{th}$ layers | 0.0066 | 0.0063 | 0.0064 |
| 156 | 5 | 32 | 0.3 | i | all layers | 0.1235 | 0.3346 | 0.2291 |
| 157 | 5 | 32 | 0.3 | ii | all layers | 0.2440 | 0.0277 | 0.1359 |
| 158 | 5 | 32 | 0.3 | iii | all layers | 0.0182 | 0.0161 | 0.0172 |
| 159 | 5 | 32 | 0.5 | i | all layers | 0.0920 | 0.2409 | 0.1665 |
| 160 | 5 | 32 | 0.5 | ii | all layers | 0.2440 | 0.0176 | 0.1308 |
| 161 | 5 | 32 | 0.5 | iii | all layers | 0.0106 | 0.0103 | 0.0104 |

**Table S3.** $R^2$-score values for the five proteins. The prediction of $g_O(\boldsymbol{r})$ was performed. In the deep-learning model, the convolution is also added to the upsampling layers.

| PDB code | All the voxels in the protein |
|---|---|
| 2ha2 | 0.988 |
| 2o4l | 0.987 |
| 3jvr | 0.987 |
| 3o5n | 0.986 |
| 4kao | 0.988 |

**Table S4.** Proteins used for the training and test (Fig. S3), different from those shown in Table 1 in the main text.

| PDB ID | Structure Title | High Reso. Limit (Å) | Dataset |
|---|---|---|---|
| 1G3D | Bovine beta-trypsin | 1.80 | Train |
| 1LBF | Indole-3-glycerol phosphate synthase | 2.05 | Train |
| 1OGD | High affinity ribose transport protein RBSD | 1.95 | Train |
| 1Q1G | Uridine phosphorylase putative | 2.02 | Train |
| 1UI0 | Uracil-DNA glycosylase | 1.50 | Train |
| 2BR1 | Chk1 kinase | 2.00 | Train |
| 2D3U | RNA polymerase from hepatitis C virus | 2.00 | Train |
| 2HA6 | Mouse acetylcholinesterase | 2.25 | Train |
| 2HB1 | Protein tyrosine phosphatase | 2.00 | Train |
| 2PK6 | HIV-1 Protease | 1.45 | Train |
| 2VWO | Factor Xa | 1.60 | Train |
| 3CCW | 3-hydroxy-3-methylglutaryl coenzyme A reductase | 2.10 | Train |

| | | | |
|---|---|---|---|
| 3FHB | Poly(ADP-ribose) polymerase | 2.30 | Train |
| 3GS6 | Glycoside hydrolase NagZ | 2.30 | Train |
| 3GY4 | Bovine trypsin | 1.55 | Train |
| 3K97 | Heat shock protein 90 | 1.95 | Train |
| 3UP2 | Aurora kinase A | 2.30 | Train |
| 4AJI | Lactate dehydrogenase A | 1.93 | Train |
| 4H3G | Beta Amyloid Cleaving Enzyme-1 | 1.85 | Train |
| 4L4Z | LsrR protein | 2.30 | Train |
| 5HVT | Human macrophage migration inhibitory factor | 1.75 | Train |
| 5L3A | Janus kinase | 1.98 | Train |
| 1UZ8 | Monoclonal antibody 291-2G3-A | 1.80 | Test |
| 2B7D | factor VIIa–tissue factor complex | 2.24 | Test |
| 3O5X | Src homology-2 domain containing protein tyrosine phosphatase-2 | 2.00 | Test |
| 4AHS | Integrase | 1.75 | Test |
| 4AZ5 | β-N-acetylhexosaminidase | 1.73 | Test |
